# The 229 chromosomes of the Atlas blue butterfly reveal rules constraining chromosome evolution in Lepidoptera

**DOI:** 10.1101/2025.03.14.642909

**Authors:** Charlotte J. Wright, Dominic Absolon, Martin Gascoigne-Pees, Roger Vila, Mara K. N. Lawniczak, Mark Blaxter

## Abstract

Chromosomal arrangements are important for processes including genetic recombination, adaptation, and speciation. Related taxa often possess similar numbers of chromosomes, but some groups show remarkable variation in chromosome numbers. Most Lepidoptera, the butterflies and moths, have 31 or 32 chromosomes, but some species deviate from this norm. We present a chromosome-level genome assembly of a heterogametic female Atlas blue butterfly (*Polyommatus atlantica*; Lycaenidae), and find it has 227 autosomes and four sex chromosomes, the highest recorded chromosome number in non-polyploid Metazoa. We show that the 227 autosomes, exceptionally small even for Lepidoptera, are derived from extensive fragmentation of the 24 ancestral lycaenid autosomes. We show that autosomal fissions likely largely occurred in euchromatic, lightly-packed regions of chromosomes. We assemble two large Z chromosomes, one of which comprises the ancestral Z fused with an autosome and retains its ancestral length, while the other is a neo-Z, formed from the fusion of an intact ancestral autosome with a fragment of a second. We find two large W chromosomes, derived from copies of the Z-linked, ancestrally autosomal sequences. In contrast to the autosomes, the sex chromosomes have not experienced fission. We observe frequent presence of chromosome-internal arrays of the telomeric repeat motif in *P. atlantica*. Such arrays are not observed in the genomes of close relatives that have not undergone fission and suggest a possible mechanism for rapid, viable fragmentation. Altogether, our findings in *P. atlantica* make evident several constraints that govern karyotypic change, a key component of eukaryotic genome evolution.

## Introduction

The rate of karyotypic evolution across the tree of life is remarkably low. Karyotypes are generally stable to such an extent that it has been possible to infer the likely ancestral linkage groups of the last common ancestor of bilaterian animals^1,2^. This stability may be because individuals that are heterozygous for inter-chromosomal rearrangements, such as the fusion or fission of a chromosome, are expected to have reduced fitness due to improper segregation during meiosis^3^. Karyotypic change may also have negative fitness consequences because chromosome rearrangements alter the folding of chromosomes and perturb patterns of recombination^4^. Consistent with this, many human cancers are associated with chromosome rearrangement^5^.

Given this conservatism, it is informative to explore the biology of groups and species where the general constraint on change appears to have been lifted. One aspect of genome structure that may modulate the frequency of interchromosomal rearrangements is centromere structure. Holocentric chromosomes, where centromeric function is dispersed across their length, are found in approximately 20% of eukaryotic species, including Lepidoptera. It has been hypothesised that holocentricity may be permissive of rapid karyotypic evolution^6,7^. In Lepidoptera, the insect order that comprises the butterflies and moths, most species have 31 chromosomes which closely correspond to the 32 ancestral linkage groups of Lepidoptera, known as Merian elements^8^. While occasional fusions are found scattered on the phylogeny, to date, high rates of fission and/or fusion are only observed in a few clades. Analysis of these clades may illuminate why karyotypes typically evolve slowly, and, because chromosomal rearrangements have been hypothesised to facilitate speciation, the processes that contribute to generation of biodiversity^9^. Heteromorphic sex chromosomes, where males and females possess cytogenetically distinct sex chromosomes, experience different evolutionary dynamics compared to autosomes. Sex chromosomes are predicted to diverge faster than autosomes (the ‘fast X effect”)^10,11^, and thus genes on sex chromosomes can play a disproportionate role in the evolution of adaptive traits and speciation. Sex chromosomes can also experience a higher rate of interchromosomal rearrangements than autosomes. For example, the heterogametic Z sex chromosome in Lepidoptera is often involved in fusions^8,12,13^.

An extreme case of chromosomal change is presented by the Atlas blue butterfly, *Polyommatus atlantica* (Elwes, 1905), where cytological examination indicated 224-226 pairs of chromosomes^14,15^. This reorganisation has been extremely rapid as species in the genus *Polyommatus*, which diverged about 3 million years ago, typically possess 23 chromosomes^16–18^. While most chromosomes in *P. atlantica* are very small, two much larger chromosomes are observed cytogenetically in males^15^. It has been suggested that sex chromosomes resist fragmentation more than autosomes^19^, raising the possibility that the large chromosomes correspond to sex chromosomes. Therefore, *P. atlantica* is an ideal species in which to understand the constraints imposed on the evolutionary trajectories of autosomes and sex chromosomes.

Here, we generate a high-quality, chromosome-level genome for *P. atlantica*. The genome consists of 227 pairs of very small autosomes derived from fission of ancestral autosomes. Two Z and two W sex chromosomes are also identified. The largest Z chromosome comprises the ancestral Z fused with an autosome while the other is a neo-Z that is a fusion of two autosomes. The two W chromosomes are derived from copies of the Z-linked sequences. Exploring the hundreds of fission events that occurred in the lineage leading to *P. atlantica*, we find that chromatin structure influences fission sites and frequent presence of internal telomeric repeat arrays. We also show that fissions do not appear to have been shaped by a requirement to retain co-regulated gene sets, but instead have occurred to generate similar sized fragments of chromosomes.

## Results

### A chromosome-level genome for *P. atlantica* consists of 229 pairs of chromosomes

To understand the evolutionary history of the *P. atlantica* genome, we generated a high-quality, chromosome-level reference genome for *P. atlantica* using PacBio HiFi data and Hi-C data derived from a single female (Figure 1A; Table 1). We assembled a 695 Mb genome (including the Z and W chromosomes), the majority of which was scaffolded into 231 scaffolds that each represent a chromosome. 69.5% of the genome was annotated as transposable elements, which is relatively high for Lepidoptera^8^. Analysis of the repeat landscape profile suggested a recent burst of LINEs and simple repeats had occurred (Figure 1D). However, the overall distribution of repeat divergence is similar to that of other closely-related lycaenid butterflies such as *P. icarus* (Rottemburg, 1775), which have a stable karyotype of *n=*23 (Figure 1D), suggesting that the dramatic change in chromosome number is not associated with a recent burst of repetitive element activity.

**Figure 1.**
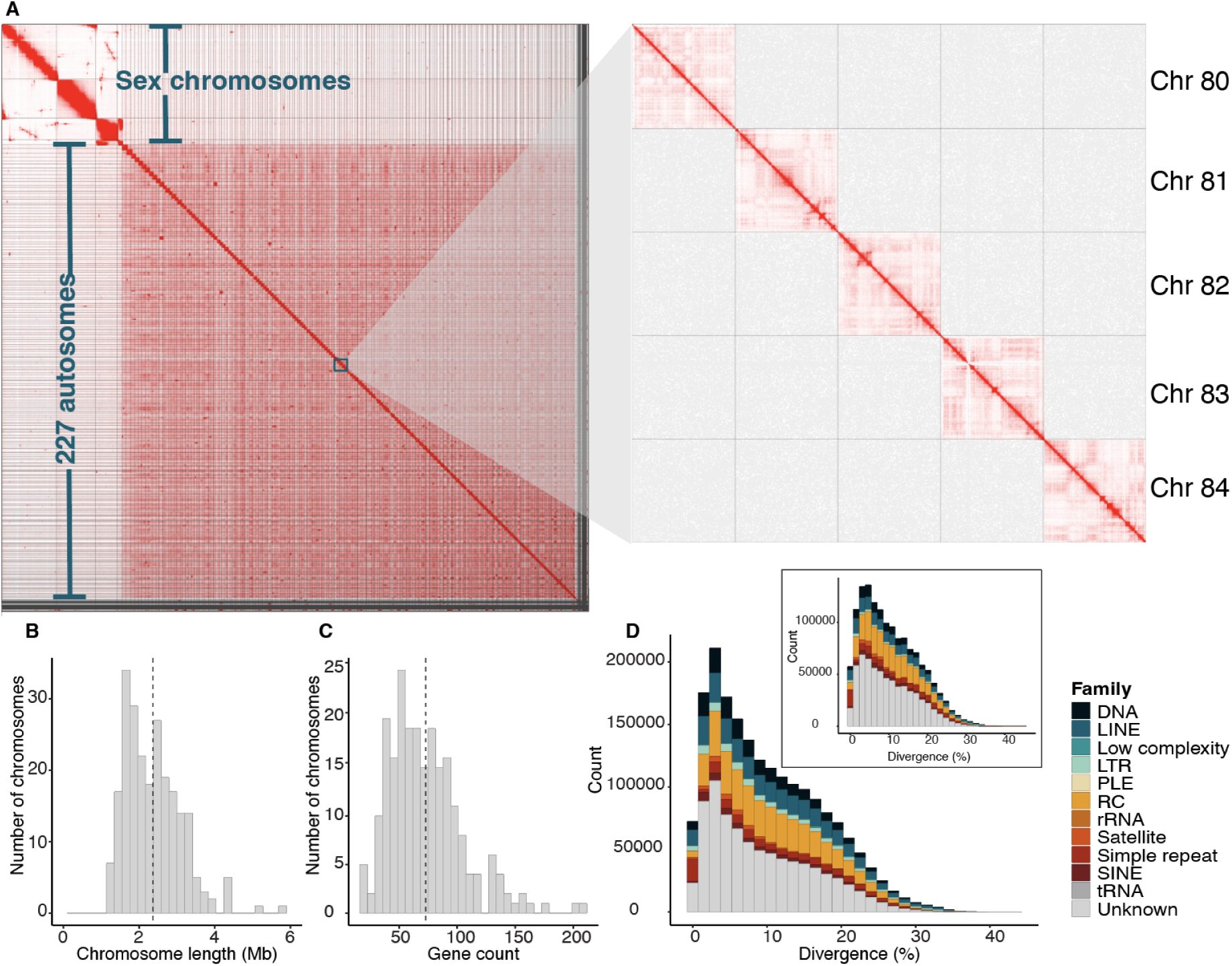
The genome of *Polyommatus atlantica* is composed of 227 autosomes of a similar size, and four large sex chromosomes. **A.** Hi-C contact map for the *P. atlantica* chromosome-level reference genome. To the right of the full map is a zoomed subset of the map for autosomes 80-84. **B.** Distribution of autosome lengths (Mb). The dashed line indicates the mean chromosome length (2.36 Mb). **C.** Distribution of the number of genes per autosome. The dashed line indicates the mean gene count (73). **D.** Repeat landscape of *P. atlantica*. Sequence divergence of each transposable element from its respective consensus sequence. Repeats are classified into repeat families. Inset in the top-right of the plot is the repeat landscape of close relative *P. icarus*.

**Table 1.** Metrics of the chromosome-level reference genomes of *Polyommatus atlantica* and *Polyommatus icarus*. * The *P. icarus* gene annotation was obtained from Ensembl. ** Genome and gene set completeness was assessed using BUSCO with the lepidoptera_odb10 dataset.

| Accession information |  |  |
| --- | --- | --- |
| Species | *Polyommatus atlantica* | *Polyommatus icarus* |
| Accession | GCA_965178725.1 | GCA_937595015.1 |
| Version | ilPolAtla1.1 | ilPollcar1.1 |
| Reference | *This study.* | Lohse *et al*, 2023 |
| Reference genome metrics |  |  |
| Assembly span (Mb) | 695.3 | 511.7 |
| Haploid chromosome number (*n*) | 229 | 23 |
| Scaffolds (*n*) | 648 | 33 |
| Scaffold N50 (Mb) | 2.7 | 21.8 |
| Span in chromosome-sized scaffolds (%) | 97.5 | 99.9 |
| Contigs | 3840 | 137 |
| Contig N50 (Mb) | 0.31 | 5.5 |
| BUSCO completeness (%) | 96.4 | 97.6 |
| BUSCO duplication (%) | 4.0 | 0.5 |
| QV | 55.4 | 57.7 |
| Feature annotation metrics |  |  |
| Genome occupied by transposable elements (%) | 69.5 | 59.6 |
| Number of protein-coding genes | 21,245 | 13,350* |
| Number of gene transcripts | 26,283 | 23,612* |
| BUSCO** completeness of gene annotation (%) | 92.7 | 96.2* |

Two Z chromosomes were identified by calculating the average read depth of PacBio HiFi data (Figure S1A; Figure S1C), and are consistent with published cytological analysis of male specimens, which identified two large chromosomes^15^. Two W chromosomes were also identified through coverage, homology to the Z chromosomes and their highly repetitive sequence content (83.0% and 71.9% for W1 and W2 respectively) (Figure S1B). The remaining 227 scaffolds had approximately twice the coverage of the Z chromosomes, consistent with being diploid autosomes. Therefore, *P. atlantica* has a chromosome complement of 227 autosomes and four distinct sex chromosomes (two Z and two W).

### Extensive fission events have resulted in 227 small autosomes of a similar size

Consistent with previous cytogenetic observations in *Polyommatus*^15,20^, there was no evidence of polyploidy in *P. atlantica*, suggesting the extreme number of chromosomes is due to many fission events. The autosomes have a mean length of 2.36 Mb with remarkably little variation (SD=0.76) (Figure 1B). Each autosome encodes an average of 73 genes (SD=33) (Figure 1C) and occupies just 0.3% (SD=0.1%) of the genome. This is much shorter than the average proportional length of chromosomes found across Lepidoptera of 3.1% per Merian element^8^. In contrast, Z1 and Z2 are 46.2 Mb and 25.5 Mb, have 1,161 and 682 genes, and occupy 6.6% and 3.7% of the genome (when including both the W and Z chromosomes), respectively.

We compared the genome of *P. atlantica* to *P. icarus* (*n*=23), which is the closest relative with a chromosome-level reference genome sequence, and found that many fissions, but no fusions, occurred in the autosomes in the lineage leading to *P. atlantica*. (Figure 2A). Each *P. atlantica* autosome was found to be homologous to part of one *P. icarus* autosome, with limited changes in gene order synteny in the fragmented chromosome compared to the intact one. Using the chromosomes of *P. icarus* can as a proxy for chromosome length in the ancestor of *P. atlantica* before fission occurred we found that the larger the ancestral chromosome, the more fissions it experienced (Spearman’s rank correlation, ⍴(21) = 0.8, *P* = 9 × 10^−6^) (Figure 2B). In Lepidoptera, smaller autosomes tend to have a higher density of repeats^8^. Consistent with the fissions having occurred recently, repeat density in the autosomes was correlated with their ancestral length before fragmentation (Spearman’s rank correlation, ⍴(235) = 0.27, *P* = 4 × 10^−5^) (Figure 2C). There was no correlation between chromosome length and repeat density in *P. atlantica* itself (Spearman’s rank correlation, ⍴(235) = −0.08, *P* =0.212) (Figure S2).

**Figure 2.**
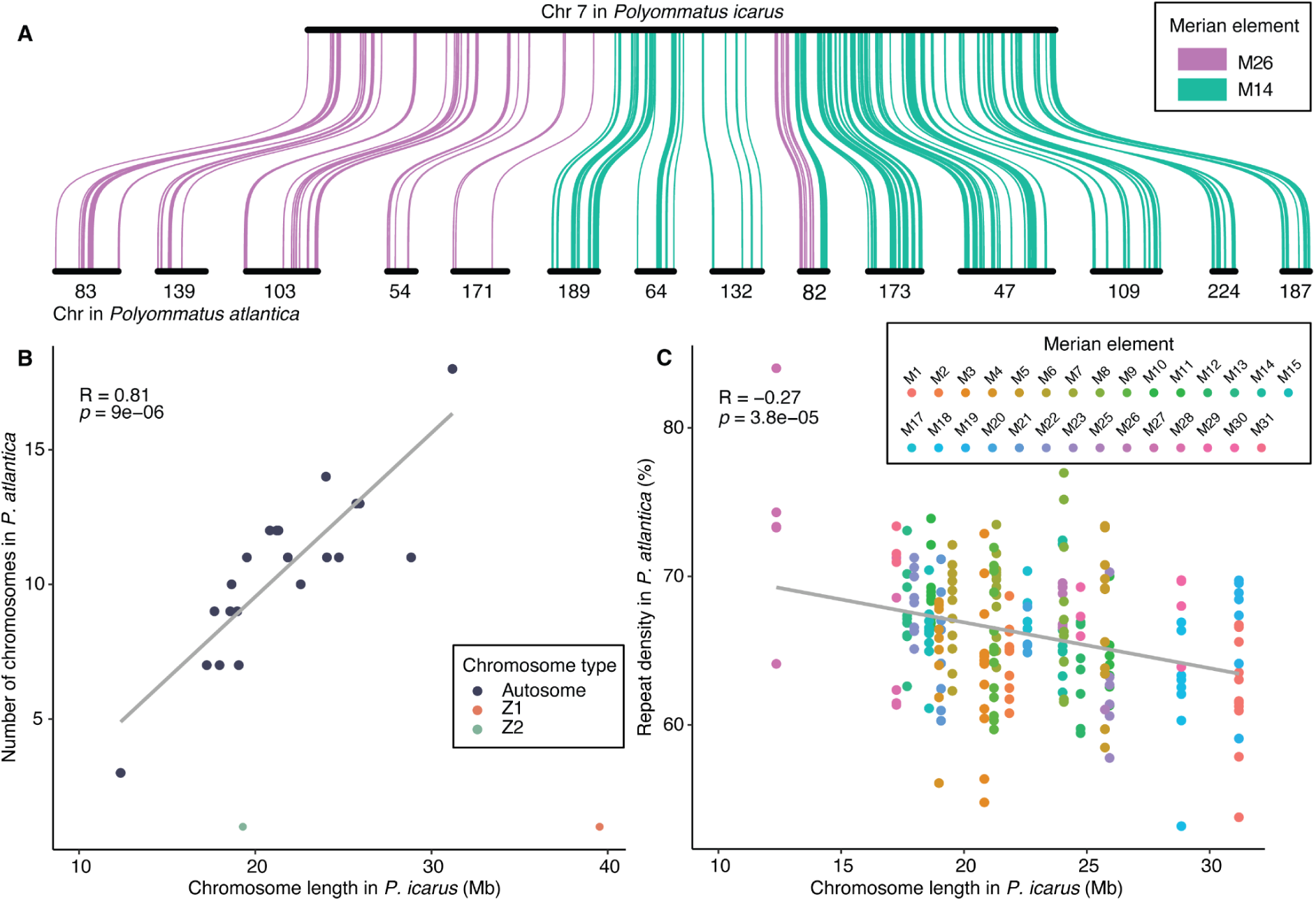
Larger chromosomes underwent more fission events, and resulting chromosomes retain sequence features associated with their length before fission. **A.** Locations of single-copy orthologues in chromosome 7 of *P. icarus* and in the 14 corresponding chromosomes of *P. atlantica*. Horizontal black bars represent chromosomes. Curved lines between species link orthologues and are coloured according to the Merian element that they belong to. Chromosome 7 of *P. icarus* is derived from an ancestral fusion of two Merian elements (M14 and M26) which was likely present in the common ancestor of *P. icarus* and *P. atlantica*. **B.** Autosome length in *P. icarus*, as a proxy for chromosome length in the last common ancestor of *Polyommatus*, is proportional to the number of chromosomes in *P. atlantica* that the chromosome has fragmented into. **C.** Autosome length in *P. icarus* is inversely correlated with repetitive sequence content (% of the chromosome occupied by repetitive elements) in the orthologous, fragmented chromosomes of *P. atlantica*. Grey lines in **B** and **C** represent the linear smoothed line through the data.

### The history of the Z chromosomes of *P. atlantica*

To investigate the origins of the sex chromosomes in *P. atlantica* (Figure 3A), each chromosome was painted with single-copy genes that define Merian elements. The largest Z chromosome (Z1) consists of MZ (the ancestral Z element) and M24 (ancestrally an autosome). The genes from these two elements have not become intermixed through intrachromosomal rearrangement (Figure 3B). The second Z chromosome (Z2) contains 95% of the M16-derived genes (125/132), suggesting that Z2 is derived from an autosome that became sex-linked. In addition, Z2 contains 10% (10/104) of the M21-derived genes, suggesting a fragment of M21 fused to this chromosome. The remaining M21-derived genes are found on 7 autosomes.

**Figure 3.**
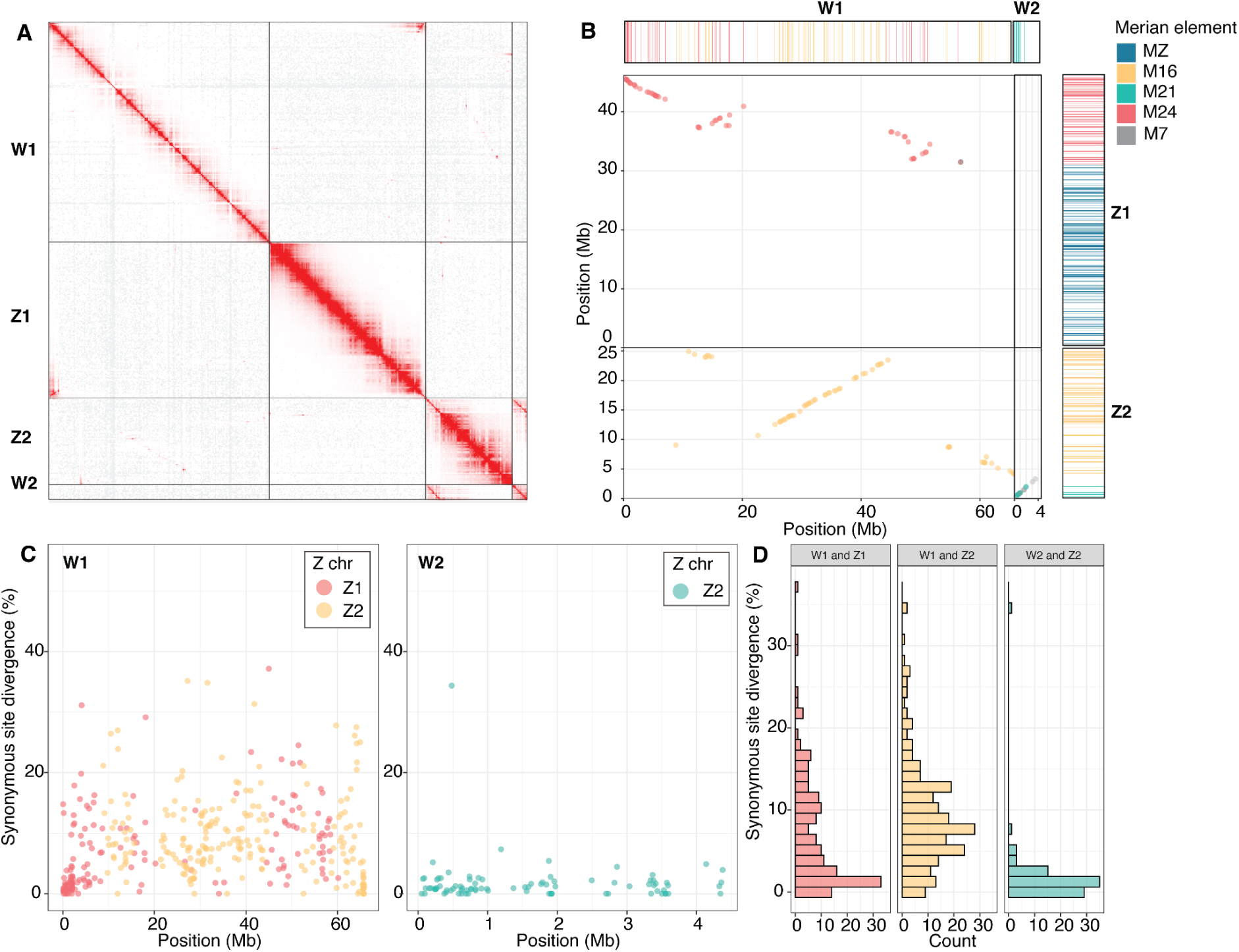
The two W chromosomes of *Polyommatus atlantica* show homology to the two Z chromosomes. **A**. Hi-C contact map of the sex chromosomes in the *P. atlantica* reference genome. **B**. Oxford plot of positions of genes in Z1 and Z2 compared to in W1 and W2, coloured by the Merian element that each gene belongs to. Rectangles above and to the right show the positions of genes used to assign membership of Merian elements. **C**. Synonymous site divergence (*d*_S_) in 464 genes in W and Z chromosomes. Points are coloured by the Z chromosome which the orthologue resides on. **D.** Distribution of *d*_S_ in genes in W and Z chromosomes.

To determine the order in which the sex chromosomes evolved, the chromosomes of *P. atlantica* were compared to those of *P. icarus*. Analysis of the coverage of PacBio HiFi read data from a female and male individual of *P. icarus* identified a single Z chromosome that corresponds to a MZ+M24 fusion (Figure S1D). This suggests the MZ+M24 fusion was present in the last common ancestor of these species. The chromosome corresponding to M16, which is Z-linked in *P. atlantica,* is autosomal in *P. icarus* (Figure S1D). This suggests that the Z2 of *P. atlantica* evolved from an autosome after its divergence from *P. icarus* approximately 3.3 million years ago^21^ and after the MZ+M24 fusion. A single autosome of *P. icarus* corresponds to M21 (Figure S1E), suggesting that a fragment of M21 joined the Z chromosome in the lineage leading to *P. atlantica.* The remaining M21-derived genes in *P. atlantica* are found on 7 autosomes.

### The W chromosomes are derived from copies of three autosomes that became sex-linked

Sex-limited chromosomes (W in ZW systems) can arise from the homolog of a chromosome that becomes Z-linked through fusion to an existing Z chromosome or by becoming a neo-Z^3,22^. Alternatively, new W sequences can arise from the fusion of an existing W chromosome to an autosome, and potentially *de novo* from a B chromosome or other material^23–25^. We identified two W chromosomes in *P. atlantica*; the first W chromosome (W1) was 65.8 Mb, making it the largest chromosome in the genome, while the second W chromosome (W2) was 4.5 Mb in length. To understand the origins and evolution of these W chromosomes, we aligned the nucleotide sequences of the W chromosomes to the other chromosomes in the genome. Alignments between W1 and Z1 covered 6.7% of W1 (Figure S3A), largely in the first 6 Mb of W1 and last 5 Mb of Z1 (Figure S3A). W1 contains 47 M24-derived genes, each of which has a corresponding copy on the M24-derived region of Z1 (Figure 3B). Therefore, this region corresponds to a previously autosomal chromosome (M24) that fused to the ancestral Z chromosome (MZ). Despite extensive degeneration of the loci on the W (it lacks 36 of the 83 M24 genes present on Z1), gene order of the retained loci has been maintained (Figure 3B). Thus, the M24 region of W1 likely evolved from the homolog of the autosome that fused to the Z chromosome.

The W1 chromosome also contains 63 M16-derived genes. These genes all have homologs on Z2, and have largely retained their gene order on W1 relative to Z2 (Figure 3B). Consistent with this, 3.6% of W1 aligned to Z2 (Figure S3A). Therefore this region of W1 may be derived from the homolog of the autosome that became Z2. W2 carries 10 M21-derived genes that have corresponding copies in the same order on Z2, and 81.9% of W2 aligned to the first 5 Mb of Z2 (Figure S3A). This suggests that W2 evolved from the homolog of the autosome (M21) that fused to a Z chromosome. We note that the nucleotide alignments and conserved genes were concentrated at one end of W2. This suggests that, while much of W2 has degenerated due to lack of recombination, recombination suppression at one end has not yet evolved or evolved relatively recently.

### Divergence estimates suggest historical recombination between Z1 and W1 chromosomes

In Lepidoptera, it is thought that Z and W chromosomes do not recombine due to achiasmatic meiosis in females^3,26^. Therefore, given that *P. icarus* has the MZ+M24 fusion, but lacks a Z2 derived from M16, we expect the M24-portions of the *P. atlantica* Z1/W1 chromosomes will be more diverged than the M16-derived regions of the Z2/W1 chromosomes. We estimated the average synonymous site divergence (*d*_S_) between orthologous regions of each pair of sex chromosomes (Figure 3C; Figure 3D). Unexpectedly, we found a significantly higher divergence between the M16-derived regions of Z2/W1 (9.4%) than between the M24-derived regions of Z1/W1 (7.2%) (Wilcoxon t-test, p < 0.001) (Figure S3B). Consistent with a fragment of M21 joining Z2 more recently, the M21-derived regions of Z2/W2 had a lower divergence (1.8%) (Wilcoxon t-test, p < 0.001).

One possible explanation for the lower divergence between the M24-derived regions of Z1/W1 than the M16-derived regions of Z2/W1, is that M16 became sex-linked earlier than expected. However, this would require a reversion of M16 back to autosomal in *P. icarus*. Alternatively, recombination may have occurred between Z1 and W1, leading to an apparently more recent divergence. Consistent with this hypothesis, the last 5 Mb of W1 was enriched for loci with under 2% divergence with Z1, compared to the rest of the W1/Z1 orthologues (Figure S4A). Analysis of gene density along the Z1 chromosome indicates that genes are evenly distributed along the chromosome, with genes with low divergence concentrated in the last 3 Mb of the chromosome (Figure S4B). The region of low divergence in W1 is spanned by many contigs, consistent with this region not being an artefact of genome assembly (Figure S4E). Moreover, in the absence of recombination, one would expect the M24-derived region of W1/Z1 in *P. icarus* and *P. atlantica* to have a consistent difference in *d*_S_ between the two species. Consistent with recombination having occurred, we found a significantly smaller difference between the *d*_S_ for genes in the first 5 Mb region of M24-derived region of W1/Z1 (Figure S4C; Figure S4D).

### Fissions occur in the A compartment and do not tend to maintain clusters of functionally-related genes

The hundreds of fissions that shaped the *P. atlantica* genome afford an excellent opportunity to understand the evolutionary constraints that influence the locations of fission events. A long-standing hypothesis is that repetitive elements facilitate chromosome rearrangements^27,28^. However, we did not find an association between sites that had undergone fission and repeat density in regions of *P. icarus* (as a proxy for repeat density in the ancestor of *P. atlantica)* that were the sites of fission (Figure S5). Fission sites may instead be related to three-dimensional genome architecture and may preferentially occur, or be permitted, in loosely-packed, transcriptionally-active, euchromatic, A compartment, instead of the repeat-rich, tightly-packed heterochromatic B compartment, or the intermediate S compartment^29^. To test this, we annotated A, B and S domains^29^ in *Cyaniris semiargus,* a close relative of *P. atlantica,* which also belongs to subtribe Polyommatina, for which dense Hi-C data were available. This provided a proxy for the ancestral chromatin landscape in the ancestor of *P. atlantica* before fission started. Consistent with our predictions, we found a significant overlap between A compartment regions and fission sites in *P. atlantica* (147/212) (p-value < 0.001, z-score = 3.621) (Figure 4A; Figure S6A) and a significant depletion in the B compartment (109/212) (p-value < 0.001, z-score = −3.747) (Figure S6B). We did not find an association between fission sites and the S compartment (56/212) (Figure S6C).

**Figure 4.**
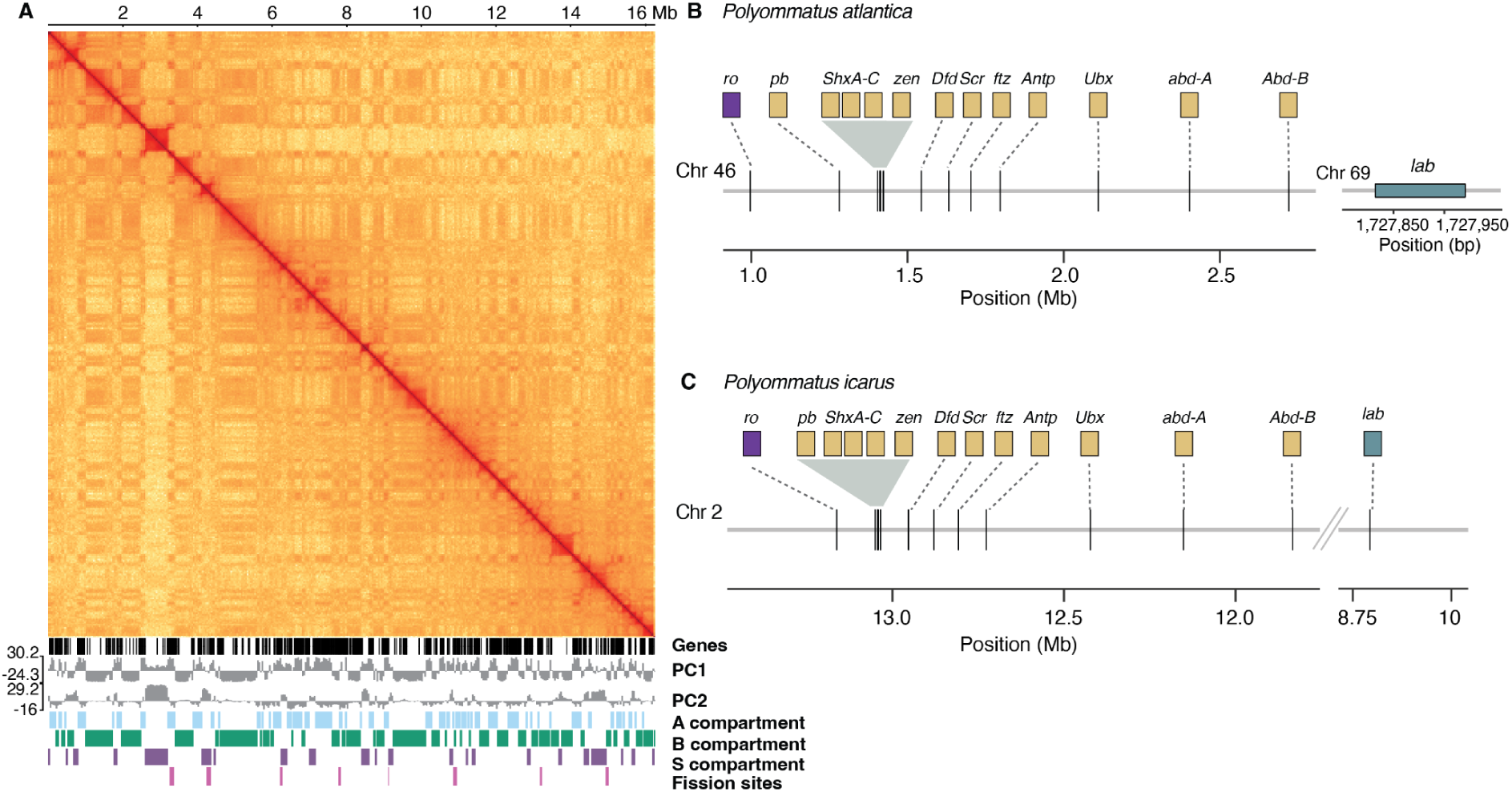
Fission sites are associated with the A compartment and depleted in the B compartment. **A**. Hi-C contact map of chromosome 14 in *C. semiargus* at 40 kb resolution. Below the matrix are gene locations, and principal components 1 (PC1) and 2 (PC2) of the Pearson’s product-moment correlation matrix. Beneath this are the locations of the three compartments (A, B and S) and locations which correspond to fissions in *P. atlantica*. **B.** Conservation of structure of the hox gene cluster in *P. atlantica*. All of the core genes in the Hox gene cluster for Lepidoptera are located in chromosome 46 in *P. atlantica* except for *lab* which is on chromosome 69. Chromosome 46 is 2.9 Mb in size and 13/56 genes on this chromosome are part of the hox gene cluster. **C,** In *P. icarus,* all genes are on chromosome 2 and are conserved in order relative to *P. atlantica*. Dashed lines in **B** and **C** indicate schematic gene locations.

Chromosomal fission could potentially disrupt long-distance *cis*-regulation of genes. Therefore, we expected that *cis*-regulated gene sets would resist fission. For example, the Hox homeobox genes have a highly constrained gene order, reflecting their complex regulation in body axis formation during development^30^. In *P. atlantica*, 9 of 10 Hox homeobox genes were located on a single chromosome (chr 46) (Table S1). The *lab* gene was found on a different chromosome (chr 69) (Figure 4B). In *P. icarus* the main cluster of nine genes spans ∼3 Mb on chromosome 2, with the *lab* locus located at the other end of the same chromosome (Figure 4C; Table S1). The preservation of the central Hox gene cluster in *P. atlantica* is consistent with a constraint on fission imposed by *cis*-regulation of gene sets. Similarly, PRD homeobox genes form a co-regulated gene cluster in Lepidoptera^30^, and this cluster was also found on a single chromosome in *P. atlantica,* suggesting a similar constraint on disruption (Table S2).

To investigate whether the co-location of homeobox genes reflects a broader constraint of co-location of related gene functions (over and above gene sets known to be *cis* co-regulated), we performed gene ontology (GO) term enrichment on each of the gene sets in the chromosomes of *P. atlantica*. We found an average of 3.1 biological processes that were significantly enriched (p<0.05) per autosome (SD=1.7). To determine whether this is higher than expected given any clustering of genes in the ancestral genome before fission occurred, we compared this enrichment score to simulated sets of fragments resulting from splitting the *P. icarus* autosomes into 227 blocks of similar size to those in the *P. atlantica* genome. This gave an average of 3.2 biological processes per block (SD=1.9), and thus no significant difference between the observed enrichment, and that obtained from random locations of fissions.

### Widespread occurrence of telomeric repeat arrays within the chromosomes of *P. atlantica*

In Lepidoptera, chromosome ends are capped by tandem repeats of the telomeric repeat sequence ([TTAGG]n). Two telomere-associated non-LTR retrotransposons, TRAS1 and SART1 insert specifically into ([TTAGG]n) repeats^31–33^. To determine whether the novel chromosome ends resulting from fission were similar to the ancestral chromosome ends, we searched for the telomeric repeat sequence. Consistent with the *de novo* addition of telomeric sequence to chromosome ends after fragmentation^20^, 54% (247/462) of the assembled chromosome ends contained at least one repeat array (Figure 5A). Chromosome ends lacking telomeric sequence are likely due to the difficulty of assembling highly repetitive subtelomeric sequence.

**Figure 5.**
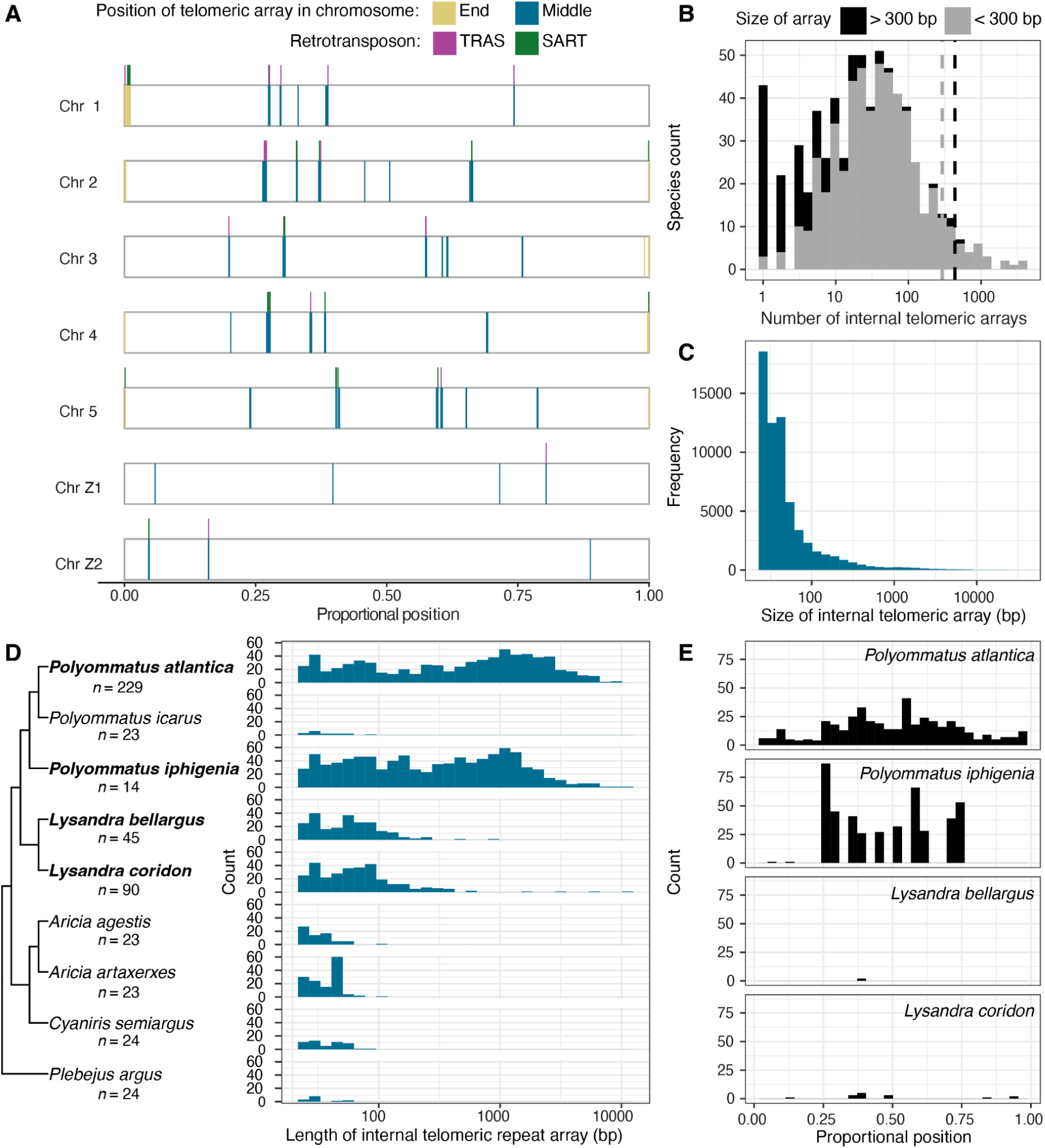
Species that have undergone many fission events have many long internal telomeric repeat arrays. **A.** Positions of telomeric repeat arrays in five autosomes and two Z chromosomes of *P. atlantica*. Each chromosome is represented by a rectangle with the positions of telomeric repeat arrays marked. Positions of TRAS and SART elements are annotated above each chromosome. Telomeric repeat arrays are coloured by whether they are located within the terminal 2.5%, or the central 95% of the chromosome. **B.** Stacked histogram of the distribution of the number of internal telomeric arrays per species, surveyed across 530 species of Lepidoptera. Each array is classified as either short (< 300 bp) or long (>=300 bp). Dashed lines indicate the values for *P. atlantica.* **C.** Distribution of the length of internal telomeric repeat arrays across 530 species of Lepidoptera. **D.** Phylogeny of Polyommatina, a subtribe within Lycaenidae, inferred from the nucleotide sequences of 4,815 single-copy orthologs. Species in bold have a haploid chromosome number that differs from the ancestral state of *n*=24 for Lycaenidae. Adjacent to each species is the length distribution of the internal telomeric arrays in the species. **E.** Distribution of the positions of long internal telomeric arrays in the chromosomes of Lycaenidae species that had at least one long array.

Unexpectedly, we found tandem telomeric repeat sequence within the main body of most chromosomes. Of the 229 chromosomes, 167 had at least one internal telomeric array (defined as arrays that occurred within the central 95% of the chromosome). Autosomes had between one (found in 46 chromosomes) to 22 arrays (found in Chr 22). Sex chromosomes also contained internal telomeric arrays (Figure 5A). The internal arrays were of the canonical telomeric repeat, with few variant copies, and frequently aIso contain TRAS and SART elements. For example, a 2 kb region of Chr 1 (3.654-3.656 Mb) consists of 260 exact copies of TTAGG and just 4 variant copies of the motif. Of the 727 internal telomeric sequences, 31% (225/727) contained TRAS and 13% contained SART (96/727). Given fusions have not occurred in *P. atlantica* and gene order is largely conserved between *P. atlantica* and *P. icarus*, the many internal telomeric arrays in *P. atlantica* do not appear to be relics of intra- or inter-chromosomal rearrangement events (Figure 3B).

The high frequency of internal telomeric sequences in *P. atlantica* led us to hypothesise that internal telomeric sequences may influence the tendency of regions to undergo fission, or modulate the extent to which fission at a given locus is deleterious. To investigate this, we searched for tandem repeats of the canonical telomeric motif in 532 Lepidopteran chromosome-level genome assemblies (Table S5). Following the convention that internal telomeric repeat arrays longer than 300 bp form a functional t-loop^34^ (hereafter referred to as long arrays), we also assessed the length distribution of internal telomeric repeat arrays. We found that 77% (214,388/277,780) of arrays were located at chromosome ends, as expected. Almost all species (530/532) contained at least one internal array (Figure 5B). However, the vast majority of internal arrays (96%; 60625/63,392) were short (<300 bp) (Figure 5B, 5C). In contrast, only 5% (28/532) of species possessed more than 10 long regions (Figure 5B).

Two species (*Chrysodeixis includens* and *Polyommatus iphigenia*) had more regions of internal telomeric sequence than *P. atlantica*, although the median length was lower in each species than in *P. atlantica*. *C. includens* is a noctuid moth that has the typical chromosome complement for a lepidopteran species (*n*=31) (Figure S7A). In contrast, *P. iphigenia* has highly rearranged chromosomes (*n*=14) that result from fusion and fission (Figure S7B). While *P. iphigenia* and *P. atlantica* both belong to Lycaenidae, karyotype data suggests that the lineages leading to *P. iphigenia* and *P. atlantica* independently underwent chromosomal change^35^. We found that species with the typical *n*=23 chromosome arrangement for Lycaenidae, including *P. icarus*, do not possess long internal telomeric arrays (Figure 5D). Consistent with previous work, we confirmed that two species of *Lysandra*, a third group within Lycaenidae that has high chromosome numbers due to fission, also have long internal telomeric arrays^36^ (15 in *L. coridon* and 2 in *L. bellargus*) (Figure 5D; Figure 5E). Thus, the presence of long internal telomeric arrays is strongly correlated with rapid chromosomal change in Lycaenidae.

### Genetic diversity does not correlate with extreme fission

Given that chromosome rearrangements have been hypothesised to be weakly deleterious and are more likely to become fixed in species with low effective population sizes^37,38^, we examined whether *P. atlantica* showed signs of recent inbreeding or low diversity relative to thirteen other species of Lycaenidae (Table S3). We found that the *P. atlantica* individual we sequenced had a genome-wide heterozygosity of 0.0043, which falls within the range of heterozygosity values exhibited by other Lycaenid individuals (0.0010-0.0069, mean 0.0029, SD 0.0016 biallelic SNPs per site) (Figure 6A).

**Figure 6.**
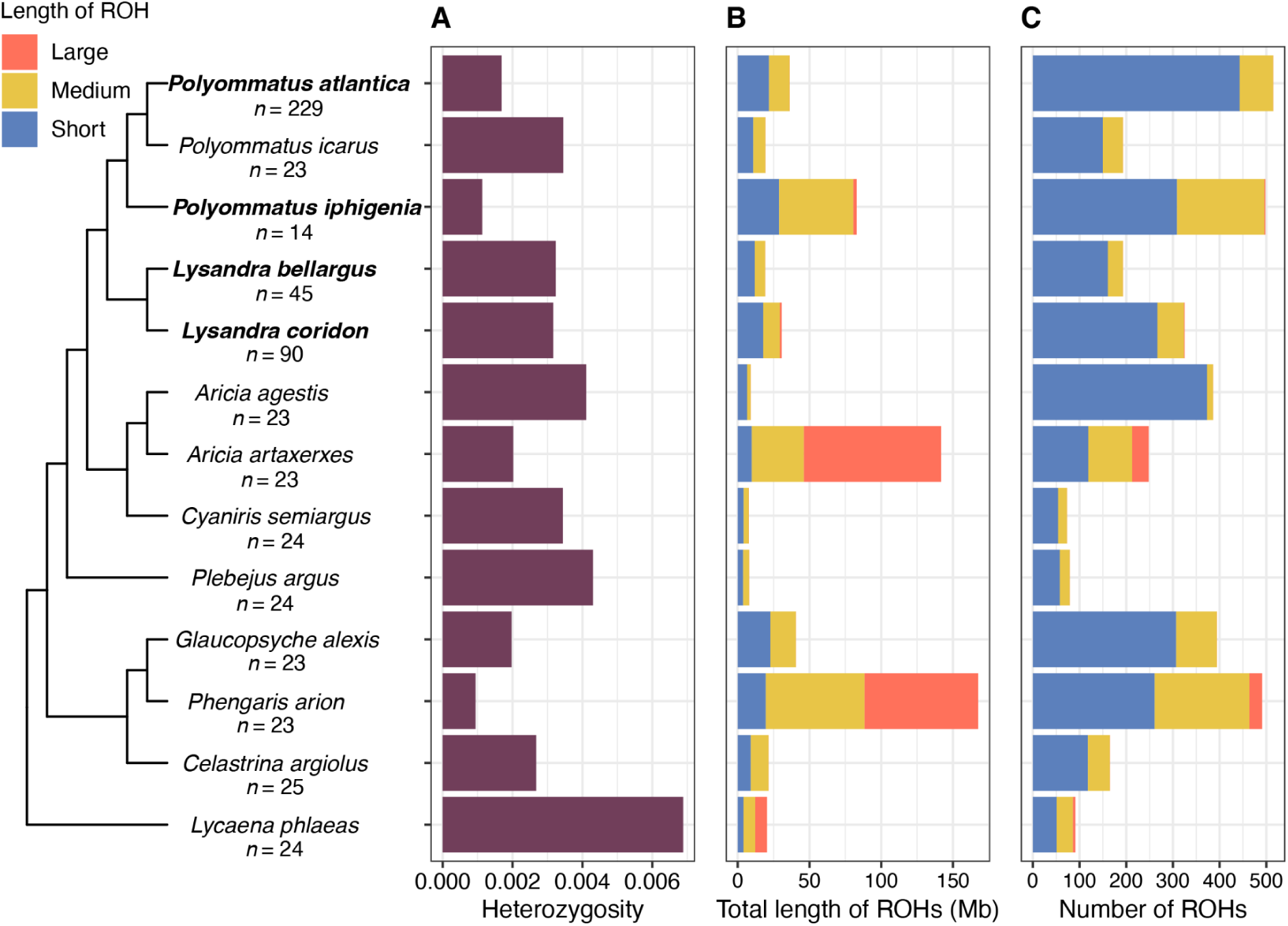
Comparable levels of heterozygosity in *P. atlantica* compared to other Lycaenids. **A.** Phylogeny of Lycaenidae inferred from the nucleotide sequences of 4,815 single-copy orthologs. Genome-wide heterozygosity (mean number of heterozygous sites per base pair, excluding sex chromosomes) per species. **B.** Total length in Mb and **C,** number of all runs of homozygosity (ROH) per species where each ROHs is classified as either short (< 100 Kb), medium (0.1 - 1 Mb) or long (> 1 Mb) in length. Species in bold have a haploid chromosome number that differs from the ancestral state of *n*=24 for Lycaenidae.

Examination of the distribution of runs of homozygosity (ROHs), which can reflect recent inbreeding, found a lower proportion of the *P. atlantica* genome is present in a ROH compared, on average, to other lycaenid individuals (7.1%) (Table S4). Previous analyses of inbreeding in Lepidoptera categorised ROHs based on their length: short ROHs (<= 0.1 Mb) were interpreted as signal of past inbreeding within small population sizes, while medium (0.1-1 Mb) indicated background relatedness and large (>=1 Mb) recent inbreeding events^39^. Following this scheme, *P. atlantica* has largely short ROHs and is devoid of long ROHs (Figure 6B; Figure 6C). However, due to the small chromosome size in *P. atlantica* (average autosome 2.2 Mb), it is unsurprising that the majority of ROHs are small.

## Discussion

*Polyommatus atlantica*, a small blue butterfly, has the highest reported diploid chromosome number in an animal that does not have a history of polyploidisation. Plants with higher numbers of chromosomes are known, but these are polyploids^40–44^. Here we show that *P. atlantica* has 229 pairs of chromosomes and that this karyotype is the product of extensive chromosome fission and not by changes in ploidy. Despite the dramatic number of fissions of autosomes, the Z and W sex chromosomes have remained intact. Fission sites were significantly associated with ancestral euchromatic regions. The *P. atlantica* genome contains many internal arrays of telomeric repeats sequences that are absent in close relatives with stable karyotypes, suggesting these arrays are associated with fragmentation.

The *P. atlantica* sex chromosomes have a complex history (Figure S8). While resolving the exact sequence of events is difficult, the sex-autosome (MZ+M24) fusion likely occurred before Z2 arose. The observed region of low divergence between the M24-derived region of Z1 and W1 is unexpected given that recombination is considered to be suppressed in lepidopteran females^26^. One explanation is that a rare recombination event occurred. In holocentric filarial nematodes that possess an X-autosome fusion, recombination between the neo-X and Y is concentrated into a short region at the distal end of the ancestrally autosomal segment due to the suppression of crossovers in the fusion-proximal region^45,46^. If a similar repatterning of recombination occurs in Lepidoptera, the observed clustering of intact genes at the end of Z1 is consistent with a similar suppression of recombination. A similar pattern of plateaus of sex-chromosome divergence was observed in a neo-Z chromosome of the butterfly *Melanargia ines*^47^. It will be informative to understand whether rare recombination is a general feature of neo-sex chromosome evolution in Lepidoptera.

Sex chromosomes often behave differently to autosomes with respect to rates of inter- and intrachromosomal rearrangements^8,48–50^. In addition to the two W chromosomes, both the ancestral Z1 chromosome and the derived Z2 chromosome have remained intact in an otherwise highly fragmented genome. The two Merian elements that are sex-linked in *P. atlantica* have undergone fission in other highly-rearranged lineages where they are autosomal^8^. This suggests that either fissions do not occur in sex chromosomes, or fissions of sex chromosomes are so deleterious that they do not reach fixation. Sex linkage may drive stability due to constraints imposed by gene dosage compensation^49^, differences in meiotic behaviour compared to autosomes, or because sex chromosomes carry genes involved in sex differentiation that need to remain in linkage^19^.

The autosomes of *P. atlantica* are similar in length. This suggests that chromosomes are required to be similar in size, and that this imposes a coarse constraint upon where fissions can occur. At a finer level, we show that chromatin structure is correlated with where fissions occur. We found that fission sites tend to have occurred in the euchromatic A compartment of the genome and not in the heterochromatic B compartment. Modelling suggests breaks occur in open chromatin in mammalian genomes^51^ and synteny breakpoints often occur in the A compartment in *Anopheles* mosquitoes^52^. Therefore, our findings suggest that architectural compartments influence genome rearrangements across monocentric and holocentric taxa. With the exception of known co-regulated clusters such as homeobox genes, we find no evidence that functionally-related clusters impose a constraint on rearrangements. In *Pieris* butterflies, which have relatively low numbers of highly rearranged chromosomes generated by fusion and fission, functional enrichment of genes in chromosomal domains has been reported^53^. One difference between the approach employed in *Pieris* and the work described here is that we account for the ancestral order of genes and thus pre-existing clustering of functionally-related genes in the ancestor. Deeper functional annotation, incorporating evidence from stage and tissue-specific transcripts, would allow possible functional clustering to be explored further.

The autosomes of *P. atlantica* include the smallest known chromosomes in Lepidoptera (excluding *P. atlantica*, the smallest is 3.3 Mb in *Leptophobia aripa*). Combined with the observation of the similar length amongst the autosomes of *P. atlantica*, this suggests that these autosomes are of the minimum size, precluding further fission. If so, each chromosome may have the minimum DNA length needed for kinetochore attachment and faithful transmission. Alternatively, each chromosome may contain a single pairing centre associated with kinetochore attachment. However, given kinetochore activity in Lepidoptera is thought to be sequence non-specific^54^, this seems less likely. Another possibility is that 2 Mb is the smallest size that permits recombination, which is needed for successful meiosis in Lepidoptera^55–57^.

The high number of internal arrays of telomeric repeats of *P. atlantica* raises the question of whether they are related to fission. The internal arrays are not derived from external telomeres due to inversions or fusions. They may be inserted during the repair of double-stranded DNA breaks^58^ or result from the activity of mobile elements. Alternatively, the internal arrays may be scars of fissions that were subsequently repaired. It is tempting to speculate that internal arrays make fissions more likely, or modulate the deleterious effect of fissions. For example, fissions at internal arrays of telomeric repeat may be more stably transmitted if these arrays provide a cap for new chromosome ends. However, while internal arrays of telomeric repeats have been implicated in chromosome rearrangements in other taxa^59^, we have not established a definitive link between these sequences and fission events in *Polyommatus*. Moreover, the conserved chromosomes of the soybean looper, *C. includens,* which has the highest number of arrays across the surveyed Lepidoptera, challenges the role of internal telomeric sequences in driving fissions. One possibility is that both number and size of the arrays may be important in determining their effects as the arrays were shorter in *C. includens* than in *P. atlantica*. It remains striking that the two lineages of *Polyommatus* with many fissions also contained numerous internal telomeric sequences. Sequencing of multiple individuals of *P. altantica* would demonstrate if there are polymorphic fissions, and if so, the relationship between these nascent fission sites and internal telomeric sequences.

*P. atlantica* is thus confirmed as having the highest known chromosome number of any diploid metazoan. All else being equal, a high number of short chromosomes should increase the rate of sequence evolution through an elevated per base recombination rate. Short term, this could be beneficial by accelerating sequence evolution which in turn enables rapid adaptation and divergence between populations. Consistent with this, fission events have been found to be associated with cladogenetic events in *Erebia* butterflies^60^. However, previous studies have observed that lineages which have undergone many chromosome rearrangements are often evolutionary young suggesting a long-term cost to highly rearranged chromosomes^8^. Therefore, while bursts of chromosomal change may drive the generation of diversity in the short-term, this may be a runaway deleterious process which leads to loss of these lineages. Exploring the relationship between the long-term consequences of such dramatic changes and speciation rates will be vital to understanding the dynamics of the generation of biodiversity.

## Methods

### Sample collection

An adult female specimen of *Polyommatus atlantica ssp. weissi* was grown from egg in the laboratory, originating from a population from the Middle Atlas mountains, Morocco (latitude 33.079, longitude −5.025). A female specimen of *Polyommatus icarus* was collected from Seva, Catalonia, Spain (latitude 41.81087, longitude 2.258244). Both specimens were snap frozen and stored at −80°C.

### DNA extraction and sequencing

Thorax tissue of *P. atlantica* was homogenised using a PowerMasher II tissue disruptor^61^. High molecular weight DNA was extracted by the Wellcome Sanger Institute Tree of Life core laboratory using a manual MagAttract protocol^62^. The DNA was sheared into 8-10 kb fragments using a g-TUBE^63^. Fragments below 8 kb were removed and the DNA was concentrated by performing a bead clean up using an automated solid-phase reversible immobilization (SPRI) with a 0.6X ratio of AMPure PB beads^64^. The concentration of the purified DNA was determined using a Nanodrop spectrophotometer and Qubit fluorometer with the Qubit dsDNA High Sensitivity Assay Kit. Fragment size distribution was assessed by running the sample on a FemtoPulse system.

The thorax tissue of *P. icarus* was homogenised using a PowerMasher II tissue disruptor^61^. High molecular weight DNA was extracted by the Wellcome Sanger Institute Tree of Life core laboratory using an automated MagAttract v2 protocol^65^. The DNA was sheared into an average fragment length of 12-20 kb using a Megaruptor 3 system^66^. Fragments shorter than 12 kb were removed by using AMPure PB beads^67^. The concentration of the purified DNA was determined using a Nanodrop spectrophotometer and Qubit fluorometer with the Qubit dsDNA High Sensitivity Assay Kit. Fragment size distribution was assessed by running the sample on the FemtoPulse system.

For *P. atlantica*, a PacBio HiFi circular consensus DNA sequencing library was prepared from the purified DNA using the Pacbio Ultra Low Input DNA Library Preparation kit and sequenced on a PacBio SEQUEL IIe flow cell by the Long Read Team of the Scientific Operations core at the Wellcome Sanger Institute. For *P. icarus*, a PacBio HiFi circular consensus DNA sequencing library was prepared from the purified DNA using the SMRTbell Prep Kit 3.0 and sequenced on a PacBio SEQUEL IIe PacBio flow cell by the Long Read Team of the Scientific Operations core at the Wellcome Sanger Institute.

The head tissue from the specimen of *P. atlantica* was used to prepare and sequence a Hi-C library by the Long Read Team of the Scientific Operations core at the Wellcome Sanger Institute. The library was prepared using the Arima Hi-C (v2) kit by following the manufacturer’s instructions and sequenced on one-eighth of an Illumina NovaSeq 6000 lane using paired-end 150 base sequencing.

### RNA extraction and sequencing

RNA was extracted from the abdomen tissue of the specimen of *P. atlantica* in the Tree of Life Laboratory at the Wellcome Sanger Institute using the RNA Extraction: Automated MagMax™ mirVana protocol^68^. RNA concentration was determined using a Nanodrop spectrophotometer and a Lunatic fluorometer using the Qubit RNA Broad-Range Assay kit. The length profile of the RNA was analysed using the Agilent RNA 6000 Pico Kit and Eukaryotic Total RNA assay. A poly(A) RNASeq library was prepared with the NEB Ultra II RNA Library Prep kit and sequenced by the Scientific Operations core at the Wellcome Sanger Institute on an Illumina NovaSeq 6000 lane using paired-end 150 base sequencing.

### Genome assembly

Genome size and heterozygosity were estimated by counting k-mers of length 31 in the PacBio HiFi reads using FastK (https://github.com/thegenemyers/FASTK) and inputting these to GenomeScope^69^. A primary assembly for *P. atlantica* was generated from the PacBio HiFi data using Hifiasm (v.0.19.8-r603)^70^ with the --primary option. Residual haplotypic duplication was removed using purge_dups (v.1.2.5)^71^. The Hi-C reads were aligned to the primary assembly using bwa-mem2 (v.2.2.1)^72^. The assembly was then scaffolded with the mapped Hi-C reads using YaHS (v.1.2a.2)^73^ using the --break option.

We screened the scaffolded assembly for scaffolds belonging to contaminant organisms by running BlobToolKit^74^ and removed contamination as described previously^75^. Three scaffolds identified as belonging to the microsporidian *Nosema bombycis* were removed. A mitochondrial genome was identified and assembled into a circular sequence with a span of 15.7 kb using MitoHifi (v.3)^76^. The nuclear assembly was assessed for biological completeness and contiguity using seqkit (v.2.8.2)^77^ and BUSCO (v.5.7.1)^78^ in miniprot mode with the lepidoptera_odb10 lineage^79^.

### Genome curation

The scaffolded assembly for *P. atlantica* was manually curated to generate a chromosome-level reference genome. The TreeVAL pipeline^80^ was implemented to prepare the necessary data required for curation including HiGlass^81^ and PretextView (https://github.com/sanger-tol/PretextView)^82^ maps and coverage and gap information. Manual curation was performed using PretextView (0.2.5, https://github.com/wtsi-hpag/PretextView) and HiGlass^81^. An initial curation effort was performed on a combined Pretext map of the primary and alternate assembly. Subsequent rounds of curation were performed on the semi-curated primary assembly to achieve the most accurate level of curation. Scaffolds were manually reordered, orientated, broken, and joined based on the Hi-C signal of support as described^75^. A total of 564 breaks and 806 joins were made and 71 false duplications were removed. This resulted in a primary nuclear assembly with a haploid chromosome number of 229 chromosomes (227 pairs of autosomes and four sex chromosomes).. This is a higher number than the *n*=224-226 previously cytologically observed^15^. However, each chromosome in the assembly was carefully inspected with detailed manual curation.

A final Hi-C contact map was generated using Juicer (v.2.0)^83^ and Juicebox (v.2.15) (available at https://github.com/aidenlab/Juicebox) (Figure 1A). A Hi-C map in Cooler file format^84^ for the final assembly was generated using bwa-mem2^85^. The base-level accuracy and k-mer completeness of the curated assembly was evaluated using Merqury (v.1.3)^86^ using k-mers of length 31 from the PacBio data.

### Gene prediction

Prior to performing gene annotation on the reference genome assembly of *P. atlantica*, a *de novo* library of transposable elements was inferred using Earl Grey (v.4.3.0)^87,88^. This repeat library was provided to RepeatMasker (v.4.1.5) to detect and soft-mask repetitive sequences^89^. Protein-coding genes and transcripts were predicted by aligning the library of short-read RNA-seq data to the soft-masked genome using HiSat2 (v.2.2.1)^90^ and providing the resulting BAM file to BRAKER (v.3.0.8)^91^ in EP mode (BRAKER2)^92^. Protein-coding genes were also independently predicted by using BRAKER in ET mode (BRAKER1)^93^, with protein hints generated using the ProtHint protein mapping pipeline (v.2.6)^94^ with the reference protein sequences for arthropods from OrthoDB ^95^ as homology evidence. The two sets of gene predictions were then combined using TSEBRA (v.1.1.2.5)^96^ with a parameter set that favoured models supported by RNA-seq evidence over those based on protein homology evidence (weight of 10,000 for RNA-seq models and 1 for protein homology models) and kept all *ab initio* models (those that did not have support from either RNA-seq or proteins). The completeness of the resulting gene set was assessed using the protein mode of BUSCO^78^ in miniprot mode with the lepidoptera_odb10 lineage^79^. Predicted protein sequences were extracted from the resulting gtf file using the script getAnnoFastaFromJoingenes from AUGUSTUS. The functions of the resulting proteins were predicted using the Pfam models in InterProScan (v.5.61-93.0). The number of protein-coding genes per chromosome was calculated using BEDtools (v.2.31.1)^97^.

### Coverage estimation

To assess per-base read coverage of each chromosome in the *P. atlantica* and *P. icarus* reference genomes, we aligned each library of PacBio reads (described above) to the respective genome using minimap2 (v.2.28-r1209)^98^. We also downloaded a PacBio library for a male individual of *P. icarus* from INSDC (run accession ERR9081703, bioproject PRJEB51265) and aligned this to the *P. icarus* reference genome. We used BEDtools (v2.31.10) to generate non-overlapping 100 kb windows and calculated coverage in each window by using mosdepth (v0.3.3)^99^.

### Butterfly genome assemblies for comparative analyses

All chromosome-level reference genomes for Lepidoptera that were on INSDC as of the 8th July 2024 were downloaded. Accession numbers for all reference genomes used are detailed in the GitHub repository https://github.com/charlottewright/Polyommatus_atlantica_MS.

### Comparison to Merian elements

Single-copy orthologues in each genome were identified using BUSCO (v.5.7.1) (miniprot mode, lepidoptera_odb10 dataset)^78^. The chromosomes of *P. atlantica* were “painted” with the positions of BUSCOs assigned to Merian elements to examine the distribution of Merian elements in the genome using lep_busco_painter (https://github.com/charlottewright/lep_busco_painter). Chromosomes were assigned to Merian elements by running lep_fusion_fission_finder (https://github.com/charlottewright/lep_fusion_fission_finder) with a window size of 3 on the orthologues. Chromosomes with fewer than three BUSCOs were separately assigned to Merian elements by determining the Merian element to which the 1 or 2 orthologues belonged to. Synteny between pairs of species was visualised using Oxford plots generated using custom scripts.

### Nucleotide alignments

Alignments between the W chromosome sequences and the other sequences in the genome of *P. atlantica* were generated with minimap2 (v.2.28-r1209)^98^, using a divergence level of below 20% (asm20) and the option -cx (intra-species asm-to-asm alignment). To remove highly repetitive regions, alignments where over 50% of the aligned region on a W chromosome was occupied by repetitive sequences were removed using custom scripts and BEDtools (v.2.31.1)^97^. Alignments were also filtered to only retain primary alignments with length >5000 bp and mapping quality >=40 using the R package pafr (https://github.com/dwinter/pafr) (v.0.0.2). To analyse the proportion of each W sequence occupied by filtered alignments, BEDtools were used to merge overlapping alignments on the W sequences.

### Divergence between the W and Z chromosomes

To estimate the average nucleotide divergence between the Z and W chromosomes of *P. atlantica*, the two W chromosomes of the *P. atlantica* reference genome were aligned to the rest of the genome using minimap2 (with the asm20 and –secondary=no parameters). Mean divergence was calculated by multiplying the gap-compressed per-base sequence divergence of each alignment by the alignment length and dividing by the total number of aligned bases.

The level of synonymous divergence (*d*_S_) between the two W chromosomes and their homologous Z chromosomes was also estimated. First, the gene annotation for *P. atlantica* (described above) was filtered to retain only the longest isoform per protein using AGAT (agat_sp_keep_longest_isoform.pl)^100^. Duplicated genes where one copy was on a Z chromosome and the other on a W chromosome were found by conducting a reciprocal best hit BLAST search^101^. In brief, the protein sequence of each Z-linked protein was compared to the rest of the proteins in the annotation (parameters: blastp -evalue 1e-5 -outfmt 5 -num_alignments 10000) (v.2.6.0)^101^. Next, the protein sequence of each W-linked protein was compared to the rest of the proteins in the annotation (parameters: blastp -evalue 1e-5-outfmt 5 -num_alignments 10000). The output was parsed to retain only W-linked proteins where the best hit was to a Z-linked protein which also had the best hit to the same W-linked protein (i.e. reciprocal best hits). Each pair of protein sequences was aligned using FSA (v.1.15.9)^102^. The alignments were then transformed into codon alignments with pal2nal (v.14)^103^. Finally, dS between each pair of sequences was estimated by using the Nei-Gojobori method implemented in codeml of PAML (v.4.10.7)^104^ with runmode=-2. Two orthologues that had dS >= 0.5 and dS <= −0.5 respectively were filtered out. Finally, mean dS was calculated per pair of W and Z chromosomes.

### Divergence between *P. atlantica* and *P. icarus*

The level of synonymous divergence between the ancestrally autosomal (M24) portion of Z1 in *P. atlantica* and Z chromosome in *P. icarus* was estimated. *P. icarus* was chosen as the closest relative to *P. atlantica* with a genome available, and a Z chromosome which has not undergone fusion as in *P. atlantica*. The chromosome-level genome was downloaded from NCBI (accession GCA_937595015.1)^105^ and a corresponding gene annotation was downloaded from Ensembl^106^. The gene annotation described above for *P. atlantica* was used. Each set of proteins was filtered with AGAT (agat_sp_keep_longest_isoform.pl) to retain only the longest isoform per protein. For *P. atlantica*, W-linked proteins were removed before running OrthoFinder (v.2.5.5)^107^ on the two protein sets to obtain 1:1 single copy orthologues between *P. atlantica* and *P. icarus.* We then selected the single copy orthogroups where the *P. icarus* protein belongs to the M24-derived portion of the *P. icarus* Z chromosome based on those where the *P. icarus* protein is located on the first 13.85 Mb of the Z chromosome (corresponding to the last location of a M24 BUSCO locus on this chromosome). Each pair of protein sequences was then aligned using FSA (v.1.15.9)^102^. The alignments were translated into codon alignments with pal2nal (v.14)^103^. The dS between each pair of sequences was estimated using the Nei-Gojobori method implemented in codeml of PAML (v.4.10.7)^104^ with runmode=-2.

### Phylogeny

Single-copy orthologues in each genome were identified using BUSCO (v.5.7.1) (using the augustus mode and the lepidoptera_odb10 dataset)^78^. The dataset was filtered using busco2fasta (available at https://github.com/lstevens17/busco2fasta) to retain the 4,815 BUSCO genes that were single-copy and present in 93% (14/15) of the genomes. The protein sequences of these BUSCO genes were aligned using MAFFT (v.7.526)^108^. Trimmed nucleotide alignments were generated by using the aligned amino acid sequences as a guide, by using trimAl (v.1.4.rev22)^109^ with parameters -gt 0.8, -st 0.001, -resoverlap 0.75, -seqoverlap 80 -backtrans. All alignments passed the alignment thresholds of trimAl. The trimmed nucleotide alignments were concatenated to form a supermatrix using catfasta2phyml (available at https://github.com/nylander/catfasta2phyml). This supermatrix was provided to IQ-TREE (v.2.03)^110^ to infer the species tree with model selection^111,112^. This chose the GTR+F+I+R9 substitution model, which specifies the GTR model with a FreeRate model with 9 rate categories^113,114^, empirical base frequencies and allowing for a proportion of invariable sites. The phylogeny was bootstrapped using 1000 ultrafast bootstrap replicates^115^.

### Telomeric repeat identification

To identify telomeric sequence, Tandem Repeats Finder (TRF) (v.4.09)^116^ was run on each genome. The output was filtered to select regions where the consensus sequence was the canonical telomeric repeat motif for Lepidoptera (TTAGG). Each region was then classified as external (i.e. at the chromosome ends) if it was located within the first or last 2.5% of the length of the chromosome, or else internal. To characterise the telomeric regions, we searched for two retrotransposon elements, TRAS and SART, which are known to be interspersed with the canonical telomeric repeat motif ^33^. Copies of these two elements were identified by searching for the sequence 5’-AAAAAAAAAACCTAACCTAACCTAACCTAA-3’ for TRAS and 5’-CCTAACCTAACCTAACCTAACCTTTTTTTTTT-3’ for SART against the reference assembly. Only hits with 100% identity to the query were retained. The co-occurrence of the canonical motif and the two elements was then found by using BEDtools intersect (v.2.31.1)^97^.

### Homeobox gene identification

Genomes were annotated with the locations of homeobox genes by using the HbxFinder pipeline from^30,117^ (https://github.com/PeterMulhair/HbxFinder). In brief, homeodomain protein sequences from three insects (*D. melanogaster, T. castaneum* and *Apis mellifera*) were used as queries in a TBLASTN (v.2.6.0)^101^ search against each genome. A single sequence was obtained per homeobox gene by filtering overlapping hits to retain the longest sequence match. These sequences were then used in a reciprocal BLASTX (v.2.6.0)^101^ search against the homeodomain protein dataset. For hits with high percentage identity (>70%), the reciprocal BLAST search enabled the initial identification of the given gene. A further round of sequence similarity searches was carried out using MMseqs2^118^ with 1 kb on either side of the homeobox genes annotated from the first BLAST search.

### Identification of breakpoints

To identify sites in *P. icarus* that were orthologous to the sites that had undergone fission in the lineage leading to *P. atlantica*, we used the locations of orthologues in the genome of *P. atlantica* and *P. icarus*. To do this, we first obtained 8,831 single copy orthologues by running OrthoFinder (v.2.5.5)^107^ on the gene annotation generated for *P. atlantica* described above and the publicly available annotation for *P. icarus* obtained from Ensembl (GCA_937595015.1, v. 2022_06) which was generated using the Ensembl Genebuild pipeline (which uses both RNA-seq and protein homology evidence)^106^. Then, a custom script (available at https://github.com/charlottewright/Polyommatus_atlantica_MS) was used to determine the locations of fissions in *P. atlantica* relative to the *P. icarus*. In brief, this compares the set of genes on each *P. atlantica* chromosome to the set of genes on the reference *P. icarus* chromosomes. The location of each gene in the query chromosomes is compared to the orthologous gene on the reference chromosomes, and thus assigned to a reference chromosome. These initial matches were filtered to only retain assignments that involve at least three genes on the reference chromosome and the query chromosome in order to minimise small regions associated with interchromosomal rearrangements. Regions along a query chromosome, where the assigned reference chromosome changes between adjacent genes, are classified as a breakpoint region. The breakpoint is taken as the end position of the coding-sequence of the last gene of a given block of synteny and the start of the first gene of the new region of synteny. The same method was also used to identify sites in *Cyaniris semiargus* (*C. semiargus*) that were orthologous to sites that had undergone fission in the lineage leading to *P. atlantica* using the gene annotation for *C. semiargus* obtained from Ensembl (GCA_905187585.1, v.2021_12) which was generated using the BRAKER2 pipeline (which uses protein homology evidence)^00^.

### Transposable element analysis

A *de novo* library of transposable elements for *P. icarus* was inferred using Earl Grey (v.4.3.0)^87,88^. The divergence of each repeat sequence from a consensus sequence was calculated by supplying the curated repeat library from Earl Grey to Repeat Masker (v.4.1.5)^89^. Enrichment for overlap of sites in *P. icarus* that were orthologous to the sites that had undergone fission in the lineage leading to *P. atlantica* with transposable elements was calculated using the overlapPermTest function implemented in regioneR (v.1.30.0)^119^ with 10,000 resamplings. In brief, this tests for the overlap between two sets of features by performing permutations while taking into account each chromosome separately. We also calculated the density of transposable elements in breakpoint regions and compared this to the density of transposable elements in *P. icarus* in 10,000 randomly generated fragments of similar size to the breakpoint regions per chromosome, using a custom script (available at https://github.com/charlottewright/Polyommatus_atlantica_MS).

### GO term analysis

A gene enrichment test was used to assess whether fission locations in *P. atlantica* have occurred in such a way as to maintain sets of genes with related biological functions. In brief, we first tested the level of gene enrichment in each autosome of *P. atlantica* and compared this level to that expected under a model of random fragmentation of autosomes in *P. icarus*. To enable comparisons between the *P. atlantica* gene sets and the simulated gene sets in *P. icarus,* all enrichment analyses were performed on single copy orthologues that were present and single copy in both genomes.

To do this, we first filtered the *P. icarus* and *P. atlantica* gene annotations to retain only the longest isoform per protein per genome using AGAT (agat_sp_keep_longest_isoform.pl)^100^. InterProScan (5.61-93.0)^120^ was used to functionally annotate each of the 8,831 single copy orthologous proteins using Pfam database and with the --goterms option to enable mapping to gene ontology (GO) terms. The resulting GO terms for each protein, along with the list of query genes, were used as input to the TopGO R package (v.2.54.0)^121^. GO term enrichment was run using the GO category ‘biological processes’ and the ‘weight01’ algorithm, which considers the GO hierarchy when calculating enrichment. Fisher’s exact test with a cut-off of p <0.05 was used to obtain significant GO terms. GO term enrichment was first performed on the sets of genes which have remained in linkage in *P. atlantica* despite fissions. As sex chromosomes have not undergone fission, only autosomes were considered for this analysis.

To account for functional clustering of genes in *P. atlantica* as a result of their co-location in the ancestral chromosomes, before fission occurred, GO term enrichment was also performed on 1,000 simulated gene sets produced by fragmenting the *P. icarus* autosomes into 227 fragments. In brief, each simulation aims to fragment the *P. icarus* autosomes into 227 autosomes by introducing 207 fissions. To fragment the autosomes in a similar way to what occurred in *P. atlantica*, we inputted a list of target autosome lengths for *P. icarus* resulting from fission. These autosome lengths were the same lengths as in the autosomes of *P. atlantica*, scaled for differences in genome size. Fissions were introduced by randomly selecting pairs of *P. icarus* chromosomes and a target fragment length. Fissions were accepted if the two fragments resulting from fission met the minimum length observed in the target fragment lengths. Fragmentation continued until 227 fragments had been produced. Finally, GO term enrichment was performed on each simulated set of *P. icarus* chromosomes.

The GO term enrichment results were summarised by calculating the mean number of enriched biological processes per fragment per simulated set of fragments. The mean number of enriched biological processes per fragment across the 1,000 simulations was then compared with the average from the observed sets of genes based on *P. atlantica*.

### Variant calling and selection of high-confidence SNPs

To estimate heterozygosity and runs of homozygosity, we performed variant calling on all genomes of Lycaenidae species for which PacBio HiFi data were available on INSDC (*n*=13) (Table S3). The variant calls for *P. atlantica* were also used to estimate divergence between the W and Z chromosomes. PacBio HiFi reads for each species of Lycaenidae were aligned to the respective genome using minimap2 (2.28-r1209) (parameters: --cs -ax map-hifi). The resulting SAM files were converted to BAM using SAMtools (v.1.6) and duplicates were marked with sambamba (v.0.6.8). Variants were called from the BAM files using DeepVariant (v.1.4.0)^122^. The VCF file was filtered to remove variants tagged as ‘RefCall’. Variants were filtered using BCFtools (v.1.17)^123^ to retain biallelic heterozygous SNPs with PHRED-quality scores >=15. Any SNPs that overlapped with annotated repeats were also filtered out to minimise erroneous calling of SNPs. As the biallelic SNPs were called from a single individual, the variant allele frequency (VAF) for each SNP is expected to be 0.5. Extreme outliers were filtered by removing SNPs with VAF<0.2 or VAF>0.8. VCF files were also filtered to retain SNPs with a minimum read coverage depth of 10 and a maximum depth of twice the mean genome-wide depth (Figure S9). The resulting VCF files contained a mean of 1,544,948 heterozygous sites (+-849,009 SD).

### Estimates of heterozygosity

The average PacBio read coverage per genome was calculated using SAMtools depth^124^ (v.1.11). Heterozygous SNP density was calculated in 10 kb windows across each genome. The total number of high-confidence heterozygous sites in each window was corrected for the total number of sites not called due to low coverage^39^. To calculate the average genome-wide heterozygosity, all 10 kb windows with sufficient coverage (defined as windows with >6,000 callable sites) were used. For species where a female was sequenced, the hemizygous W and Z chromosomes were excluded from this calculation.

### Identification of runs of homozygosity

Runs of homozygosity (ROH) were defined as regions showing at least 25% lower heterozygosity than expected based on the genome-wide average. Average heterozygosity per 100 kb region was calculated by averaging the heterozygosity from the 10 constituent 10 kb bins, as used in Bortoluzzi *et al*^39^. Only 10 kb bins with sufficient coverage (defined at bins with >6,000 callable sites) were included. If the heterozygosity value of the 100 kb bin was below 25% of the average genome-wide heterozygosity value, the 100 kb bin was inferred to contain a ROH. The inferred ROHs were categorised by length: small (<= 0.1 Mb), medium (0.1-1 Mb) and long (>= 1 Mb), following Bortoluzzi *et al*^39^. These categories represent past relatedness, background relatedness and recent inbreeding/relatedness respectively.

### Hi-C processing and compartment calling

Hi-C data for *Cyaniris semiargus* was downloaded from NCBI and aligned to the *C. semiargus* genome (accession GCA_905187585.1)^125^ using bwa (v.0.7.17-r1188)^85^. *C. semiargus* was chosen for this due to the availability of Hi-C data for this species and its close relatedness to *P. atlantica.* A corresponding gene annotation was downloaded from Ensembl^106^. A Pairtools (v.0.3.0)^126^ was used to parse the alignments into Hi-C pairs (with –add-columns mapq) and sort the pairs. Duplicates were detected and removed using pairtools dedup.

Architectural compartments were called by following a pipeline similar to that developed for *Bombyx mori*^29^. This detects three compartments (A, B and S). A Hi-C contact matrix was then generated at 40 kb resolution using the cloud pairix function of the cooler suite (v.0.9.3)^84^. The matrix was then balanced (normalised) using the balance function in cooler. Finally, the zoomify function of the cooler suite was used to generate matrices at multiple resolutions. The Hi-C matrix at 40 kb resolution was converted to a sparse matrix using SciPy (v.1.5.3)^127^. The sparse matrix was transformed into an observed/expected intra-chromosomal matrix using the “obs_exp_matrix_non_zero” function supplied by HiCExplorer (v.3.7.2)^128^, then into the Pearson’s product-moment correlation matrix with the function “np.corrcoef”. The resulting matrix was then normalised by scaling to unit variance by using the ‘standard scaler’ function of Scikit learn (v.0.23.2)^129^. Next, the first three principal components were fitted using the PCA function of Scikit learn. A k-means clustering strategy was then applied to the principal component vector by using the “Kmeans” function of Scikit learn and a cluster size of 3. These three clusters were assigned as the A, B and S compartment. Each 40 kb bin was then assigned to a cluster and thus compartment based on the cluster to which it was closest to. The average size of the annotated compartments in *C. semiargus* were 134 kb, 215 kb and 106 kb for A, B and S respectively. A total percentage of 29.9%, 45.8% and 14.4% of the *C. semiargus* genome was assigned to A, B and S respectively. Enrichment for overlap of breakpoints with compartment type was calculated using the R package regioneR (v.1.30.0)^119^ which performs a permutation test (10,000 times) while taking into account each chromosome separately. Feature locations were plotted together with the Hi-C data using pyGenomeTracks (v3.9)^130^

## Data availability

The reference genome of *Polyommatus atlantica* generated in this study has been deposited in ENA under the BioProject accession PRJEB86335. The PacBio data for *Polyommatus icarus* generated in this study has been deposited in ENA under the BioProject accession PRJEB51265. The reference genomes analysed in this study are available at NCBI and the accession numbers are detailed in Table S5. Large data files associated with this paper, including repeat annotations, repeat libraries and VCF files are available at the Zenodo repository https://doi.org/10.5281/zenodo.15014702131. The code and data associated with the analyses and figures can be found in the GitHub repository (https://github.com/charlottewright/Polyommatus_atlantica_MS).

## Supplemental information

Document S1. Figures S1–S9 and Tables S1-S4

Table S2. Excel file containing additional data too large to fit in a PDF, related to Table 5

## Acknowledgements

C.J.W., M.B. and M.K.N.L. acknowledge funding support from the Wellcome Trust award 220540/Z/20/A ‘Wellcome Sanger Institute Quinquennial Review 2021–2026’. M.B. and M.K.N.L. acknowledge funding support from Wellcome Trust grant no. 218328. R.V. is supported by grant PID2022-139689NB-I00 (MICIU/ AEI/ 10.13039/501100011033 and ERDF, EU) and by grant 2021-SGR-00420 (Departament de Recerca i Universitats, Generalitat de Catalunya). We would like to express our thanks for the expert support from the Sanger Scientific Operations core and the Sanger Tree of Life Core Laboratory for DNA extraction and generating sequencing data. We would also like to thank Héloïse Muller for technical advice on calling architectural compartments, and Lewis Stevens for advice on analyses and for providing feedback on the manuscript. Thanks also to Peter Holland and Ian Henderson for providing feedback on the manuscript. For the purpose of Open Access, the author has applied a CC BY public copyright licence to any Author Accepted Manuscript version arising from this submission.

## Contributions

C. J. W. and M. B. conceptualised the project. R. V. and M. G-P. collected the specimens. D. A. manually curated the reference genome, and M.K.N.L and M. B supervised the project. C. J. W. analysed the data and drafted the manuscript. All authors commented on, revised, and approved the manuscript.

## Reference

1. Putnam, N.H., Butts, T., Ferrier, D.E.K., Furlong, R.F., Hellsten, U., Kawashima, T., Robinson-Rechavi, M., Shoguchi, E., Terry, A., Yu, J.-K., et al. (2008). The amphioxus genome and the evolution of the chordate karyotype. Nature 453, 1064–1071.

2. Simakov, O., Bredeson, J., Berkoff, K., Marletaz, F., Mitros, T., Schultz, D.T., O’Connell, B.L., Dear, P., Martinez, D.E., Steele, R.E., et al. (2022). Deeply conserved synteny and the evolution of metazoan chromosomes. Sci Adv 8, eabi5884.

3. White, M.J.D. (1973). Animal Cytology and Evolution, 3rd edn Cambridge University Press (Cambridge University Press).

4. Vara, C., Paytuví-Gallart, A., Cuartero, Y., Álvarez-González, L., Marín-Gual, L., Garcia, F., Florit-Sabater, B., Capilla, L., Sanchéz-Guillén, R.A., Sarrate, Z., et al. (2021). The impact of chromosomal fusions on 3D genome folding and recombination in the germ line. Nat. Commun. 12, 2981.

5. Leibowitz, M.L., Zhang, C.-Z., and Pellman, D. (2015). Chromothripsis: A new mechanism for rapid karyotype evolution. Annu. Rev. Genet. 49, 183–211.

6. Lucek, K., Augustijnen, H., and Escudero, M. (2022). A holocentric twist to chromosomal speciation? Trends Ecol. Evol. 37, 655–662.

7. Senaratne, A.P., Cortes-Silva, N., and Drinnenberg, I.A. (2022). Evolution of holocentric chromosomes: Drivers, diversity, and deterrents. Semin. Cell Dev. Biol. 127, 90–99.

8. Wright, C.J., Stevens, L., Mackintosh, A., Lawniczak, M., and Blaxter, M. (2024). Comparative genomics reveals the dynamics of chromosome evolution in Lepidoptera. Nat Ecol Evol 8, 777–790.

9. Faria, R., and Navarro, A. (2010). Chromosomal speciation revisited: rearranging theory with pieces of evidence. Trends Ecol. Evol. 25, 660–669.

10. Vicoso, B., and Charlesworth, B. (2006). Evolution on the X chromosome: unusual patterns and processes. Nat. Rev. Genet. 7, 645–653.

11. Charlesworth, B., Coyne, J.A., and Barton, N.H. (1987). The Relative Rates of Evolution of Sex Chromosomes and Autosomes. Am. Nat. 130, 113–146.

12. Mongue, A.J., Nguyen, P., Voleníková, A., and Walters, J.R. (2017). Neo-sex Chromosomes in the Monarch Butterfly, Danaus plexippus. G3 7, 3281–3294.

13. Nguyen, P., Sýkorová, M., Šíchová, J., Kůta, V., Dalíková, M., Čapková Frydrychová, R., Neven, L.G., Sahara, K., and Marec, F. (2013). Neo-sex chromosomes and adaptive potential in tortricid pests. Proc. Natl. Acad. Sci. U. S. A. 110, 6931–6936.

14. de Lesse, H. (1960). Speciation et variation chromosomique chez les Lepidopteres Rhopaloceres. Annales des sciences naturelles. Zoologie et biologie animale, 1–223.

15. Lukhtanov, V.A. (2015). The blue butterfly Polyommatus (Plebicula) atlanticus (Lepidoptera, Lycaenidae) holds the record of the highest number of chromosomes in the non-polyploid eukaryotic organisms. Comp. Cytogenet. 9, 683–690.

16. Robinson, R. (1971). Lepidoptera Genetics (Oxford: Pergamon Press).

17. Makino, S. (1951). An atlas of the chromosome numbers in animals. An atlas of the chromosome numbers in animals.

18. Talavera, G., Lukhtanov, V., Pierce, N.E., and Vila, R. (2022). DNA barcodes combined with multilocus data of representative taxa can generate reliable higher-level phylogenies. Syst. Biol. 71, 382–395.

19. White, M.J.D. (1946). The evidence against polyploidy in sexually-reproducing animals. Am. Nat. 80, 610–618.

20. Hospodářská, M., Voleníková, A.C., Koutecký, P., Vila, R., Talavera, G., Provazníková, I., Dalíková, M., and Nguyen, P. (2024). Polyommatine blue butterflies reveal unexpected integrity of the W sex chromosome amid extensive chromosome fragmentation. bioRxiv, 2024.06.25.600692. 10.1101/2024.06.25.600692.

21. Talavera, G., Lukhtanov, V.A., Pierce, N.E., and Vila, R. (2013). Establishing criteria for higher-level classification using molecular data: the systematics of Polyommatus blue butterflies (Lepidoptera, Lycaenidae). Cladistics 29, 166–192.

22. Rice, W.R. (1996). Evolution of the Y sex chromosome in animals. Bioscience 46, 331–343.

23. Charlesworth, D., Charlesworth, B., and Marais, G. (2005). Steps in the evolution of heteromorphic sex chromosomes. Heredity (Edinb.) 95, 118–128.

24. Bachtrog, D. (2013). Y-chromosome evolution: emerging insights into processes of Y-chromosome degeneration. Nat. Rev. Genet. 14, 113–124.

25. Fraïsse, C., Picard, M.A.L., and Vicoso, B. (2017). The deep conservation of the Lepidoptera Z chromosome suggests a non-canonical origin of the W. Nat. Commun. 8, 1486.

26. Turner, J.R., and Sheppard, P.M. (1975). Absence of crossing-over in female butterflies (Heliconius). Heredity 34 *Part 2*, 265–269.

27. Montgomery, E.A., Huang, S.M., Langley, C.H., and Judd, B.H. (1991). Chromosome rearrangement by ectopic recombination in Drosophila melanogaster: genome structure and evolution. Genetics 129, 1085–1098.

28. Mieczkowski, P.A., Lemoine, F.J., and Petes, T.D. (2006). Recombination between retrotransposons as a source of chromosome rearrangements in the yeast Saccharomyces cerevisiae. DNA Repair 5, 1010–1020.

29. Gil, J., Rosin, L.F., Navarrete, E., Chowdhury, N., Abraham, S., Cornilleau, G., Lei, E.P., Mozziconacci, J., Mirny, L.A., Muller, H., et al. (2023). Unique territorial and sub-chromosomal organization revealed in the holocentric moth Bombyx mori. bioRxiv. 10.1101/2023.09.14.557757.

30. Mulhair, P.O., Crowley, L., Boyes, D.H., Harper, A., Lewis, O.T., Darwin Tree of Life Consortium, and Holland, P.W.H. (2023). Diversity, duplication, and genomic organization of homeobox genes in Lepidoptera. Genome Res. 33, 32–44.

31. Okazaki, S., Ishikawa, H., and Fujiwara, H. (1995). Structural analysis of TRAS1, a novel family of telomeric repeat-associated retrotransposons in the silkworm, Bombyx mori. Mol. Cell. Biol. 15, 4545–4552.

32. Takahashi, H., Okazaki, S., and Fujiwara, H. (1997). A new family of site-specific retrotransposons, SART1, is inserted into telomeric repeats of the silkworm, Bombyx mori. Nucleic Acids Res. 25, 1578–1584.

33. Lukhtanov, V.A., and Pazhenkova, E.A. (2023). Diversity and evolution of telomeric motifs and telomere DNA organization in insects. Biol. J. Linn. Soc. Lond. 140, 536–555.

34. Watson, J.M., Trieb, J., Troestl, M., Renfrew, K., Mandakova, T., Fulnecek, J., Shippen, D.E., and Riha, K. (2021). A hypomorphic allele of telomerase uncovers the minimal functional length of telomeres in Arabidopsis. Genetics 219. 10.1093/genetics/iyab126.

35. Kandul, N.P., Lukhtanov, V.A., Dantchenko, A.V., Coleman, J.W.S., Sekercioglu, C.H., Haig, D., and Pierce, N.E. (2004). Phylogeny of Agrodiaetus Hübner 1822 (Lepidoptera: Lycaenidae) inferred from mtDNA sequences of COI and COII and nuclear sequences of EF1-alpha: karyotype diversification and species radiation. Syst. Biol. 53, 278–298.

36. Pazhenkova, E.A., and Lukhtanov, V.A. (2023). Chromosomal conservatism vs chromosomal megaevolution: enigma of karyotypic evolution in Lepidoptera. Chromosome Res. 31, 16.

37. Bush, G.L., Case, S.M., Wilson, A.C., and Patton, J.L. (1977). Rapid speciation and chromosomal evolution in mammals. Proc. Natl. Acad. Sci. U. S. A. 74, 3942–3946.

38. Wilson, A.C., Bush, G.L., Case, S.M., and King, M.C. (1975). Social structuring of mammalian populations and rate of chromosomal evolution. Proc. Natl. Acad. Sci. U. S. A. 72, 5061–5065.

39. Bortoluzzi, C., Wright, C.J., Lee, S., Cousins, T., Genez, T.A.L., Thybert, D., Martin, F.J., Haggerty, L., The Darwin Tree of Life Project Consortium, Blaxter, M., et al. (2023). Lepidoptera genomics based on 88 chromosomal reference sequences informs population genetic parameters for conservation. bioRxiv, 2023.04.14.536868. 10.1101/2023.04.14.536868.

40. Röser, M. (2015). Mitosis and interphase of the highly polyploid palm voanioalagerardii (2n = 606 ± 3). Cytogenet. Genome Res. 147, 70–79.

41. Oginuma, K., Munzinger, J., and Tobe, H. (2006). Exceedingly high chromosome number in Strasburgeriaceae, a monotypic family endemic to New Caledonia. Osterr. Bot. Z. 262, 97–101.

42. Love, A. (1977). Cytotaxonomical atlas of the Pteridophyta. J. Camer, Vaduz.

43. Uhl, C.H. (1978). Chromosomes of Mexican Sedum II. Section pachysedum. Rhodora 80, 491–512.

44. Guo, Z.-Y., Liu, H.-M., Wang, K.-K., Fujiwara, T., Liu, Z.-Y., Zhang, X.-C., and Schneider, H. (2024). Huperziacrassifolia (Lycopodiaceae), a new species from China based on morphological characters and molecular evidence. PhytoKeys 246, 27–42.

45. Stevens, L., Kieninger, M., Chan, B., Wood, J.M.D., Gonzalez de la Rosa, P., Allen, J., and Blaxter, M. (2024). The genome of Litomosoides sigmodontis illuminates the origins of Y chromosomes in filarial nematodes. PLoS Genet. 20, e1011116.

46. Henzel, J.V., Nabeshima, K., Schvarzstein, M., Turner, B.E., Villeneuve, A.M., and Hillers, K.J. (2011). An asymmetric chromosome pair undergoes synaptic adjustment and crossover redistribution during Caenorhabditis elegans meiosis: implications for sex chromosome evolution. Genetics 187, 685–699.

47. Decroly, T., Vila, R., Lohse, K., and Mackintosh, A. (2024). Rewinding the ratchet: rare recombination locally rescues neo-W degeneration and generates plateaus of sex-chromosome divergence. Mol. Biol. Evol. 10.1093/molbev/msae124.

48. Mathers, T.C., Wouters, R.H.M., Mugford, S.T., Swarbreck, D., van Oosterhout, C., and Hogenhout, S.A. (2021). Chromosome-Scale Genome Assemblies of Aphids Reveal Extensively Rearranged Autosomes and Long-Term Conservation of the X Chromosome. Mol. Biol. Evol. 38, 856–875.

49. Ohno, S. (2013). Sex Chromosomes and Sex-Linked Genes (Springer Science & Business Media).

50. Ahola, V., Lehtonen, R., Somervuo, P., Salmela, L., Koskinen, P., Rastas, P., Välimäki, N., Paulin, L., Kvist, J., Wahlberg, N., et al. (2014). The Glanville fritillary genome retains an ancient karyotype and reveals selective chromosomal fusions in Lepidoptera. Nat. Commun. 5, 4737.

51. Berthelot, C., Muffato, M., Abecassis, J., and Roest Crollius, H. (2015). The 3D organization of chromatin explains evolutionary fragile genomic regions. Cell Rep. 10, 1913–1924.

52. Lukyanchikova, V., Nuriddinov, M., Belokopytova, P., Taskina, A., Liang, J., Reijnders, M.J.M.F., Ruzzante, L., Feron, R., Waterhouse, R.M., Wu, Y., et al. (2022). Anopheles mosquitoes reveal new principles of 3D genome organization in insects. Nat. Commun. 13, 1960.

53. Hill, J., Rastas, P., Hornett, E.A., Neethiraj, R., Clark, N., Morehouse, N., de la Paz Celorio-Mancera, M., Cols, J.C., Dircksen, H., Meslin, C., et al. (2019). Unprecedented reorganization of holocentric chromosomes provides insights into the enigma of lepidopteran chromosome evolution. Sci Adv 5, eaau3648.

54. Senaratne, A.P., Muller, H., Fryer, K.A., Kawamoto, M., Katsuma, S., and Drinnenberg, I.A. (2021). Formation of the CenH3-Deficient Holocentromere in Lepidoptera Avoids Active Chromatin. Curr. Biol. 31, 173–181.e7.

55. Yasukochi, Y., Ashakumary, L.A., Baba, K., Yoshido, A., and Sahara, K. (2006). A second-generation integrated map of the silkworm reveals synteny and conserved gene order between lepidopteran insects. Genetics 173, 1319–1328.

56. Jiggins, C.D., Mavarez, J., Beltrán, M., McMillan, W.O., Johnston, J.S., and Bermingham, E. (2005). A genetic linkage map of the mimetic butterfly Heliconius melpomene. Genetics 171, 557–570.

57. Kaback, D.B., Guacci, V., Barber, D., and Mahon, J.W. (1992). Chromosome size-dependent control of meiotic recombination. Science 256, 228–232.

58. Nergadze, S.G., Santagostino, M.A., Salzano, A., Mondello, C., and Giulotto, E. (2007). Contribution of telomerase RNA retrotranscription to DNA double-strand break repair during mammalian genome evolution. Genome Biol. 8, R260.

59. Aksenova, A.Y., Greenwell, P.W., Dominska, M., Shishkin, A.A., Kim, J.C., Petes, T.D., and Mirkin, S.M. (2013). Genome rearrangements caused by interstitial telomeric sequences in yeast. Proc. Natl. Acad. Sci. U. S. A. 110, 19866–19871.

60. Augustijnen, H., Bätscher, L., Cesanek, M., Chkhartishvili, T., Dincă, V., Iankoshvili, G., Ogawa, K., Vila, R., Klopfstein, S., de Vos, J.M., et al. (2024). A macroevolutionary role for chromosomal fusion and fission in Erebia butterflies. Sci. Adv. 10, eadl0989.

61. Denton, A., Oatley, G., Cornwell, C., Quail, M., and Howard, C. (2023). Sanger Tree of Life Sample Homogenisation: PowerMash. protocols.io.

62. Strickland, M., Moll, R., Cornwell, C., Smith, M., and Howard, C. (2023). Sanger Tree of Life HMW DNA Extraction: Manual MagAttract.

63. Oatley, G., Sampaio, F., Kitchin, L., do Amaral, R.J.V., and Howard, C. (2023). Sanger Tree of Life HMW DNA Fragmentation: Covaris g-TUBE for ULI PacBio. protocols.io. 10.17504/protocols.io.q26g7pm81gwz/v1.

64. Oatley, G., Sampaio, F., and Howard, C. (2023). Sanger Tree of Life Fragmented DNA clean up: Automated SPRI. protocols.io.

65. Oatley, G., Denton, A., and Howard, C. (2023). Sanger Tree of Life HMW DNA Extraction: Automated MagAttract v.2. protocols.io. 10.17504/protocols.io.kxygx3y4dg8j/v1.

66. Bates, A., Clayton-Lucey, I., and Howard, C. (2023). Sanger Tree of Life HMW DNA Fragmentation: Diagenode Megaruptor®3 for LI PacBio. protocols.io. 10.17504/protocols.io.81wgbxzq3lpk/v1.

67. Strickland, M., Cornwell, C., and Howard, C. (2023). Sanger Tree of Life Fragmented DNA clean up: Manual SPRI. protocols.io. 10.17504/protocols.io.kxygx3y1dg8j/v1.

68. do Amaral, R.J.V., Bates, A.A.B., Denton, A., Yatsenko, H., Jay, J., and Howard, C. (2023). Sanger Tree of Life RNA Extraction: Automated MagMax^TM^ mirVana. protocols.io.

69. Ranallo-Benavidez, T.R., Jaron, K.S., and Schatz, M.C. (2020). GenomeScope 2.0 and Smudgeplot for reference-free profiling of polyploid genomes. Nat. Commun. 11, 1432.

70. Cheng, H., Concepcion, G.T., Feng, X., Zhang, H., and Li, H. (2021). Haplotype-resolved de novo assembly using phased assembly graphs with hifiasm. Nat. Methods 18, 170–175.

71. Guan, D., McCarthy, S.A., Wood, J., Howe, K., Wang, Y., and Durbin, R. (2020). Identifying and removing haplotypic duplication in primary genome assemblies. Bioinformatics 36, 2896–2898.

72. Li, H. (2013). Aligning sequence reads, clone sequences and assembly contigs with BWA-MEM. arXiv [q-bio.GN].

73. Zhou, C., McCarthy, S.A., and Durbin, R. (2023). YaHS: yet another Hi-C scaffolding tool. Bioinformatics 39. 10.1093/bioinformatics/btac808.

74. Challis, R., Richards, E., Rajan, J., Cochrane, G., and Blaxter, M. (2020). BlobToolKit - Interactive Quality Assessment of Genome Assemblies. G3 10, 1361–1374.

75. Howe, K., Chow, W., Collins, J., Pelan, S., Pointon, D.-L., Sims, Y., Torrance, J., Tracey, A., and Wood, J. (2021). Significantly improving the quality of genome assemblies through curation. Gigascience 10. 10.1093/gigascience/giaa153.

76. Uliano-Silva, M., Ferreira, J.G.R.N., Krasheninnikova, K., Darwin Tree of Life Consortium, Formenti, G., Abueg, L., Torrance, J., Myers, E.W., Durbin, R., Blaxter, M., et al. (2023). MitoHiFi: a python pipeline for mitochondrial genome assembly from PacBio high fidelity reads. BMC Bioinformatics 24, 288.

77. Shen, W., Le, S., Li, Y., and Hu, F. (2016). SeqKit: A Cross-Platform and Ultrafast Toolkit for FASTA/Q File Manipulation. PLoS One 11, e0163962.

78. Simão, F.A., Waterhouse, R.M., Ioannidis, P., Kriventseva, E.V., and Zdobnov, E.M. (2015). BUSCO: assessing genome assembly and annotation completeness with single-copy orthologs. Bioinformatics 31, 3210–3212.

79. Kriventseva, E.V., Kuznetsov, D., Tegenfeldt, F., Manni, M., Dias, R., Simão, F.A., and Zdobnov, E.M. (2019). OrthoDB v10: sampling the diversity of animal, plant, fungal, protist, bacterial and viral genomes for evolutionary and functional annotations of orthologs. Nucleic Acids Res. 47, D807–D811.

80. Pointon Damon-Lee, Eagles Will, Sims Yumi, Muffato Matthieu, Surana Priyanka, Qi Guoying sanger-tol/treeval: v1.1.1 - Ancient Aurora (H1) 10.5281/zenodo.12653164.

81. Kerpedjiev, P., Abdennur, N., Lekschas, F., McCallum, C., Dinkla, K., Strobelt, H., Luber, J.M., Ouellette, S.B., Azhir, A., Kumar, N., et al. (2018). HiGlass: web-based visual exploration and analysis of genome interaction maps. Genome Biol. 19, 125.

82. Harry, E. (2022). PretextView (Paired REad TEXTure Viewer): A desktop application for viewing pretext contact maps.

83. Durand, N.C., Shamim, M.S., Machol, I., Rao, S.S.P., Huntley, M.H., Lander, E.S., and Aiden, E.L. (2016). Juicer Provides a One-Click System for Analyzing Loop-Resolution Hi-C Experiments. Cell Syst 3, 95–98.

84. Abdennur, N., and Mirny, L.A. (2020). Cooler: scalable storage for Hi-C data and other genomically labeled arrays. Bioinformatics 36, 311–316.

85. Vasimuddin, M., Misra, S., Li, H., and Aluru, S. (2019). Efficient Architecture-Aware Acceleration of BWA-MEM for Multicore Systems. In 2019 IEEE International Parallel and Distributed Processing Symposium (IPDPS) (IEEE), pp. 314–324.

86. Rhie, A., Walenz, B.P., Koren, S., and Phillippy, A.M. (2020). Merqury: reference-free quality, completeness, and phasing assessment for genome assemblies. Genome Biol. 21, 245.

87. Baril, T., Imrie, R.M., and Hayward, A. (2022). Earl Grey: a fully automated user-friendly transposable element annotation and analysis pipeline. 10.21203/rs.3.rs-1812599/v1.

88. Baril, T., Imrie, R., and Hayward, A. (2021). TobyBaril/EarlGrey: Earl Grey v1.2 10.5281/zenodo.5718734.

89. Smit, A., Hubley, R., and Green, P. (2015). Repeatmasker open-4.0. http://www.repeatmasker.org.

90. Kim, D., Paggi, J.M., Park, C., Bennett, C., and Salzberg, S.L. (2019). Graph-based genome alignment and genotyping with HISAT2 and HISAT-genotype. Nat. Biotechnol. 37, 907–915.

91. Gabriel, L., Brůna, T., Hoff, K.J., Ebel, M., Lomsadze, A., Borodovsky, M., and Stanke, M. (2024). BRAKER3: Fully automated genome annotation using RNA-seq and protein evidence with GeneMark-ETP, AUGUSTUS, and TSEBRA. Genome Res. 34, 769–777.

92. Brůna, T., Hoff, K.J., Lomsadze, A., Stanke, M., and Borodovsky, M. (2021). BRAKER2: automatic eukaryotic genome annotation with GeneMark-EP+ and AUGUSTUS supported by a protein database. NAR Genom Bioinform 3, lqaa108.

93. Hoff, K.J., Lange, S., Lomsadze, A., Borodovsky, M., and Stanke, M. (2016). BRAKER1: Unsupervised RNA-Seq-Based Genome Annotation with GeneMark-ET and AUGUSTUS. Bioinformatics 32, 767–769.

94. Brůna, T., Lomsadze, A., and Borodovsky, M. (2020). GeneMark-EP+: eukaryotic gene prediction with self-training in the space of genes and proteins. NAR Genom Bioinform 2, lqaa026.

95. Kuznetsov, D., Tegenfeldt, F., Manni, M., Seppey, M., Berkeley, M., Kriventseva, E.V., and Zdobnov, E.M. (2023). OrthoDB v11: annotation of orthologs in the widest sampling of organismal diversity. Nucleic Acids Res. 51, D445–D451.

96. Gabriel, L., Hoff, K.J., Brůna, T., Borodovsky, M., and Stanke, M. (2021). TSEBRA: transcript selector for BRAKER. BMC Bioinformatics 22, 566.

97. Quinlan, A.R., and Hall, I.M. (2010). BEDTools: a flexible suite of utilities for comparing genomic features. Bioinformatics 26, 841–842.

98. Li, H. (2018). Minimap2: pairwise alignment for nucleotide sequences. Bioinformatics 34, 3094–3100.

99. Pedersen, B.S., and Quinlan, A.R. (2018). Mosdepth: quick coverage calculation for genomes and exomes. Bioinformatics 34, 867–868.

100. Dainat, J., Hereñú, D., Davis, E., Crouch, K., LucileSol, Agostinho, N., pascal-git, and tayyrov (2022). NBISweden/AGAT: AGAT-v1.0.0 10.5281/zenodo.7255559.

101. Camacho, C., Coulouris, G., Avagyan, V., Ma, N., Papadopoulos, J., Bealer, K., and Madden, T.L. (2009). BLAST+: architecture and applications. BMC Bioinformatics 10, 421.

102. Bradley, R.K., Roberts, A., Smoot, M., Juvekar, S., Do, J., Dewey, C., Holmes, I., and Pachter, L. (2009). Fast statistical alignment. PLoS Comput. Biol. 5, e1000392.

103. Suyama, M., Torrents, D., and Bork, P. (2006). PAL2NAL: robust conversion of protein sequence alignments into the corresponding codon alignments. Nucleic Acids Res. 34, W609–W612.

104. Yang, Z. (2007). PAML 4: phylogenetic analysis by maximum likelihood. Mol. Biol. Evol. 24, 1586–1591.

105. Lohse, K., Wellcome Sanger Institute Tree of Life programme, Wellcome Sanger Institute Scientific Operations: DNA Pipelines collective, Tree of Life Core Informatics collective, and Darwin Tree of Life Consortium (2023). The genome sequence of the Common Blue, Polyommatus icarus (Rottemburg, 1775) [version 1; peer review: awaiting peer review]. Wellcome Open Res. 8, 72.

106. Cunningham, F., Allen, J.E., Allen, J., Alvarez-Jarreta, J., Amode, M.R., Armean, I.M., Austine-Orimoloye, O., Azov, A.G., Barnes, I., Bennett, R., et al. (2022). Ensembl 2022. Nucleic Acids Res. 50, D988–D995.

107. Emms, D.M., and Kelly, S. (2019). OrthoFinder: phylogenetic orthology inference for comparative genomics. Genome Biol. 20, 238.

108. Katoh, K., and Standley, D.M. (2013). MAFFT multiple sequence alignment software version 7: improvements in performance and usability. Mol. Biol. Evol. 30, 772–780.

109. Capella-Gutiérrez, S., Silla-Martínez, J.M., and Gabaldón, T. (2009). trimAl: a tool for automated alignment trimming in large-scale phylogenetic analyses. Bioinformatics 25, 1972–1973.

110. Nguyen, L.-T., Schmidt, H.A., von Haeseler, A., and Minh, B.Q. (2015). IQ-TREE: a fast and effective stochastic algorithm for estimating maximum-likelihood phylogenies. Mol. Biol. Evol. 32, 268–274.

111. Kalyaanamoorthy, S., Minh, B.Q., Wong, T.K.F., von Haeseler, A., and Jermiin, L.S. (2017). ModelFinder: fast model selection for accurate phylogenetic estimates. Nat. Methods 14, 587–589.

112. Chernomor, O., Von, H.A., and Minh, B.Q. (2016). Terrace aware data structure for phylogenomic inference from supermatrices. Systematic biology 65, 997–1008.

113. Yang, Z. (1995). A space-time process model for the evolution of DNA sequences. Genetics 139, 993–1005.

114. Soubrier, J., Steel, M., Lee, M.S.Y., Der Sarkissian, C., Guindon, S., Ho, S.Y.W., and Cooper, A. (2012). The influence of rate heterogeneity among sites on the time dependence of molecular rates. Mol. Biol. Evol. 29, 3345–3358.

115. Hoang, D.T., Chernomor, O., von Haeseler, A., Minh, B.Q., and Vinh, L.S. (2018). UFBoot2: Improving the Ultrafast Bootstrap Approximation. Mol. Biol. Evol. 35, 518–522.

116. Benson, G. (1999). Tandem repeats finder: a program to analyze DNA sequences. Nucleic Acids Res. 27, 573–580.

117. Zhong, Y.-F., and Holland, P.W.H. (2011). HomeoDB2: functional expansion of a comparative homeobox gene database for evolutionary developmental biology. Evol. Dev. 13, 567–568.

118. Steinegger, M., and Söding, J. (2017). MMseqs2 enables sensitive protein sequence searching for the analysis of massive data sets. Nat. Biotechnol. 35, 1026–1028.

119. Gel, B., Díez-Villanueva, A., Serra, E., Buschbeck, M., Peinado, M.A., and Malinverni, R. (2016). regioneR: an R/Bioconductor package for the association analysis of genomic regions based on permutation tests. Bioinformatics 32, 289–291.

120. Jones, P., Binns, D., Chang, H.-Y., Fraser, M., Li, W., McAnulla, C., McWilliam, H., Maslen, J., Mitchell, A., Nuka, G., et al. (2014). InterProScan 5: genome-scale protein function classification. Bioinformatics 30, 1236–1240.

121. Alexa, A., and Rahnenfuhrer, J. (2010). topGO: enrichment analysis for gene ontology. R package version.

122. Poplin, R., Chang, P.-C., Alexander, D., Schwartz, S., Colthurst, T., Ku, A., Newburger, D., Dijamco, J., Nguyen, N., Afshar, P.T., et al. (2018). A universal SNP and small-indel variant caller using deep neural networks. Nat. Biotechnol. 36, 983–987.

123. Danecek, P., Bonfield, J.K., Liddle, J., Marshall, J., Ohan, V., Pollard, M.O., Whitwham, A., Keane, T., McCarthy, S.A., Davies, R.M., et al. (2021). Twelve years of SAMtools and BCFtools. Gigascience 10. 10.1093/gigascience/giab008.

124. Li, H., Handsaker, B., Wysoker, A., Fennell, T., Ruan, J., Homer, N., Marth, G., Abecasis, G., Durbin, R., and 1000 Genome Project Data Processing Subgroup (2009). The Sequence Alignment/Map format and SAMtools. Bioinformatics 25, 2078–2079.

125. Lohse, K., Hayward, A., Laetsch, D.R., Marques, V., Vila, R., Tyler-Smith, C., Wellcome Sanger Institute Tree of Life programme, Wellcome Sanger Institute Scientific Operations: DNA Pipelines collective, Tree of Life Core Informatics collective, and Darwin Tree of Life Consortium (2023). The genome sequence of the Mazarine Blue, Cyaniris semiargus (Rottemburg, 1775). Wellcome Open Res. 8, 181.

126. Open2C, Abdennur, N., Fudenberg, G., Flyamer, I.M., Galitsyna, A.A., Goloborodko, A., Imakaev, M., and Venev, S.V. (2024). Pairtools: From sequencing data to chromosome contacts. PLoS Comput. Biol. 20, e1012164.

127. Virtanen, P., Gommers, R., Oliphant, T.E., Haberland, M., Reddy, T., Cournapeau, D., and Van Mulbregt P. (2020). SciPy 1.0: fundamental algorithms for scientific computing in Python. Nature methods 17, 261–272.

128. Ramírez, F., Bhardwaj, V., Arrigoni, L., Lam, K.C., Grüning, B.A., Villaveces, J., Habermann, B., Akhtar, A., and Manke, T. (2018). High-resolution TADs reveal DNA sequences underlying genome organization in flies. Nat. Commun. 9, 189.

129. Pedregosa, F., Varoquaux, G., Gramfort, A., Michel, V., Thirion, B., Grisel, O., and Duchesnay (2011). Scikit-learn: Machine learning in Python. the Journal of machine Learning research 12, 2825–2830.

130. Lopez-Delisle, L., Rabbani, L., Wolff, J., Bhardwaj, V., Backofen, R., Grüning, B., Ramírez, F., and Manke, T. (2021). pyGenomeTracks: reproducible plots for multivariate genomic datasets. Bioinformatics 37, 422–423.

131. Wright, C.J. (2025). The 229 chromosomes of the Atlas blue butterfly reveal rules constraining chromosome evolution in Lepidoptera. 10.5281/zenodo.15014702.

