## Supplementary_information for "The 229 chromosomes of the Atlas blue butterfly reveal rules constraining chromosome evolution in Lepidoptera"

### Contents


Table S1. Locations of Hox genes in *Polyommatus atlantica* and *Polyommatus icarus*. 2

Table S2. Locations of PRD genes in *Polyommatus atlantica*. 3

Table S3. Details of genomes and PacBio data HiFi used for variant calling. 4

Table S4. Genome-wide heterozygosity and prevalence of runs of homozygosity across Lycaenidae 5

Figure S1. Identification of sex-linked scaffolds in the *P. atlantica* and *P. icarus* reference assemblies reveals *P. atlantic*a has two Z chromosomes while *P. icarus* has a single Z chromosome. 7

Figure S2. Autosome length is not correlated with average repeat density in *P. atlantica*. 8

Figure S3. Nucleotide alignments of W1 and W2 to Z1 and Z2 in *P. atlantica* and distribution of the average synonymous site divergence per pair of sex chromosome 9

Figure S4. A region of low synonymous site divergence on Z1 and W1. 10

Figure S5. Fission sites of *P. atlantica* are not associated with highly repetitive regions 12

Figure S6. Correlation between the ancestral chromatin landscape and fission sites in *P. atlantica*. 13

Figure S7. Merian element paints of *Polyommatus iphigenia* and *Chrysodeixis includen*s. 14

Figure S8. Models of potential routes by which the Z1Z2/W1W2 chromosome system in *P. atlantica* evolved. 15

Figure S9. Quality control and filtering of variants. 17

| Species | Chr | Start | Stop | Hox gene | Orientation | Percent identity score |
|---|---|---|---|---|---|---|
| *P. atlantica* | SUPER_46 | 996157 | 996313 | Ro | L | 70 |
| *P. atlantica* | SUPER_46 | 1281560 | 1281680 | Pb | L | 100 |
| *P. atlantica* | SUPER_46 | 1404254 | 1404431 | ShxA | L | 65 |
| *P. atlantica* | SUPER_46 | 1411276 | 1411453 | ShxB | R | 56 |
| *P. atlantica* | SUPER_46 | 1413236 | 1413416 | ShxC | L | 65 |
| *P. atlantica* | SUPER_46 | 1422820 | 1423003 | zen | L | 80 |
| *P. atlantica* | SUPER_46 | 1544158 | 1544335 | Dfd | R | 100 |
| *P. atlantica* | SUPER_46 | 1632013 | 1632199 | Scr | L | 100 |
| *P. atlantica* | SUPER_46 | 1702234 | 1702411 | ftz | L | 82 |
| *P. atlantica* | SUPER_46 | 1796048 | 1796225 | Antp | L | 100 |
| *P. atlantica* | SUPER_46 | 2110501 | 2110684 | Ubx | L | 100 |
| *P. atlantica* | SUPER_46 | 2402702 | 2402885 | abd-A | L | 100 |
| *P. atlantica* | SUPER_46 | 2720573 | 2720750 | Abd-B | L | 100 |
| *P. atlantica* | SUPER_69 | 1727814 | 1727991 | lab | L | 98 |
| *P. icarus* | OW569322.1 | 1012739 | 1012916 | lab | R | 98.305 |
| *P. icarus* | OW569322.1 | 11832845 | 11833022 | Abd-B | R | 100 |
| *P. icarus* | OW569322.1 | 12151501 | 12151684 | abd-A | R | 100 |
| *P. icarus* | OW569322.1 | 12422919 | 12423102 | Ubx | R | 100 |
| *P. icarus* | OW569322.1 | 12726340 | 12726517 | Antp | R | 100 |
| *P. icarus* | OW569322.1 | 12807226 | 12807403 | ftz | R | 82.609 |
| *P. icarus* | OW569322.1 | 12879042 | 12879228 | Scr | R | 100 |
| *P. icarus* | OW569322.1 | 12952999 | 12953176 | Dfd | L | 100 |
| *P. icarus* | OW569322.1 | 13034595 | 13034778 | zen | R | 79.31 |
| *P. icarus* | OW569322.1 | 13041294 | 13041474 | ShxC | R | 66.102 |
| *P. icarus* | OW569322.1 | 13043285 | 13043462 | ShxB | L | 56.14 |
| *P. icarus* | OW569322.1 | 13050514 | 13050691 | ShxA | R | 66.102 |
| *P. icarus* | OW569322.1 | 13162148 | 13162343 | Pb | R | 100 |
| *P. icarus* | OW569322.1 | 13402788 | 13402920 | Ro | R | 75 |

**Table S1. Locations of Hox genes in *Polyommatus atlantica* and *Polyommatus icarus*.**

| Chromosome | Start (bp) | Stop (bp) | PRD gene | Orientation | Percent identity score |
|---|---|---|---|---|---|
| SUPER_193 | 431738 | 431780 | Pax3/7 | L | 87.5 |
| SUPER_193 | 432187 | 432334 | Pax3/7 | L | 87.5 |
| SUPER_193 | 527872 | 528022 | Pax3/7 | R | 93.75 |
| SUPER_193 | 528415 | 528463 | Pax3/7 | R | 93.75 |
| SUPER_83 | 77239 | 77413 | Hbn | R | 91.489 |
| SUPER_83 | 97275 | 97452 | Arx | R | 69.565 |
| SUPER_83 | 282188 | 282239 | Rax | L | 100 |
| SUPER_83 | 320527 | 320671 | Rax | L | 100 |
| SUPER_83 | 475276 | 475417 | Otp | L | 85.417 |
| SUPER_145 | 698052 | 698226 | Phox | R | 100 |
| SUPER_145 | 724284 | 724368 | Phox | R | 100 |
| SUPER_1 | 1079013 | 1079151 | Prrx | R | 89.362 |
| SUPER_1 | 1080707 | 1080746 | Prrx | R | 100 |
| SUPER_83 | 77239 | 77413 | Hbn | R | 91.489 |
| SUPER_83 | 97275 | 97452 | Arx | R | 69.565 |
| SUPER_83 | 282188 | 282239 | Rax | L | 100 |
| SUPER_83 | 320527 | 320671 | Rax | L | 100 |
| SUPER_83 | 475276 | 475417 | Otp | L | 85.417 |

**Table S2. Locations of PRD genes in *Polyommatus atlantica.***

| Species | Accession number for genome | Bioproject | SRA run for PacBio data | Reference |
|---|---|---|---|---|
| *Aricia agestis* | GCA_905147365.1 | PRJEB41901 | ERR6544653 | (Hayward et al. 2023)[1] |
| *Aricia artaxerxes* | GCA_937612035.1 | PRJEB52239 | ERR9439488 | (Ebdon *et al.* 2022)[2] |
| *Celastrina argiolus* | GCA_905187575.2 | PRJEB42086 | ERR6558180 | (Hayward *et al*, 2021)[3] |
| *Cyaniris semiargus* | GCA_905187585.1 | PRJEB42089 | ERR6560795 | (Lohse *et al*, 2022)[4] |
| *Glaucopsyche alexis* | GCA_905404095.1 | PRJEB43781 | ERR6436366 | (Hinojosa Galisteo *et al*, 2021)[5] |
| *Lycaena phlaeas* | GCA_905333005.2 | PRJEB43522 | ERR6576321 | (Lohse *et al*, 2021)[6] |
| *Lysandra bellargus* | GCA_905333045.1 | PRJEB43523 | ERR6576322 | (Lohse *et al*, 2022)[7] |
| *Lysandra coridon* | GCA_905220515.1 | PRJEB42939 | ERR6576318 | (Vila *et al*, 2023)[8] |
| *Phengaris arion* | GCA_963565745.1 | PRJEB65385 | ERR11892476 | (Meredith *et al*, 2024) [9] |
| *Plebejus argus* | GCA_905404155.3 | PRJEB43788 | ERR6606789 | (Hayward *et al*, 2022)[10] |
| *Polyommatus atlantica* | *This study* | PRJEB50432 | ERR12736855, ERR10753933 | *This study* |
| *Polyommatus icarus* | GCA_937595015.1 | PRJEB51265 | ERR9081703 | (Lohse *et al*, 2023)[11] |
| *Polyommatus iphigenia* | GCA_963422495.1 | PRJEB64749 | ERR11809156 | |

**Table S3. Details of genomes and PacBio data HiFi used for variant calling.**

Variant calling was performed on all genomes of Lycaenidae species for which PacBio HiFi data were available on INSDC as of July 2024.

| Species | Max depth | Genome size (Mb) | Number of heterozygous sites | Genome-wide heterozygosity | No. of short ROHs | No. of medium ROHs | No. of long ROHs | Sum of short ROHs (Mb) | Sum of medium ROHs (Mb) | Sum of long ROHs (Mb) | Genome occupied by ROHs (%) |
|---|---|---|---|---|---|---|---|---|---|---|---|
| *Aricia agestis* | 102 | 435.3 | 3558094 | 0.004209 | 373 | 13 | 0 | 6.6 | 2.37 | 0 | 2.06 |
| *Aricia artaxerxes* | 53 | 458.5 | 921881 | 0.002089 | 119 | 94 | 36 | 9.87 | 36.24 | 96.66 | 31.14 |
| *Celastrina argiolus* | 59 | 499.1 | 1319215 | 0.00286 | 118 | 47 | 0 | 9.05 | 12.39 | 0 | 4.30 |
| *Cyaniris semiargus* | 85 | 441.5 | 1509656 | 0.003498 | 54 | 19 | 0 | 4.1 | 3.59 | 0 | 1.74 |
| *Glaucopsyche alexis* | 77 | 619.5 | 1192391 | 0.001984 | 307 | 87 | 0 | 22.74 | 17.8 | 0 | 6.54 |
| *Lycaena phlaeas* | 107 | 420.5 | 2875123 | 0.006892 | 51 | 35 | 5 | 4.03 | 8.27 | 8.13 | 4.86 |
| *Lysandra bellargus* | 104 | 528.9 | 1593978 | 0.003271 | 161 | 32 | 0 | 11.9 | 7.17 | 0 | 3.61 |
| *Lysandra coridon* | 93 | 540.7 | 1695282 | 0.003239 | 267 | 57 | 1 | 17.78 | 11.66 | 1.15 | 5.656 |
| *Phengaris arion* | 286 | 544.5 | 483875 | 0.000951 | 261 | 203 | 29 | 19.6 | 68.92 | 81.08 | 31.15 |
| *Plebejus argus* | 43 | 382.1 | 1494047 | 0.004301 | 58 | 21 | 0 | 3.78 | 4.21 | 0 | 2.09 |
| *Polyommatus atlantica* | 68 | 658.3 | 1074732 | 0.002117 | 443 | 72 | 0 | 21.82 | 14.31 | 0 | 5.49 |
| *Polyommatus icarus* | 66 | 511.8 | 1742675 | 0.003518 | 150 | 43 | 0 | 10.9 | 8.37 | 0 | 3.77 |
| *Polyommatus iphigenia* | 89 | 39.3 | 623381 | 0.001181 | 309 | 187 | 3 | 28.86 | 51.7 | 3.26 | 213.15 |

**Table S4. Genome-wide heterozygosity and prevalence of runs of homozygosity across Lycaenidae**

The SNPs in each genome were filtered to retain SNPs with a minimum depth of 10 and a maximum depth (max depth in the table) of twice the mean genome-wide depth. The genome-wide heterozygosity (mean number of heterozygous sites per base pair, excluding sex chromosomes) and prevalence of runs of homozygosity (ROHs) in each sampled species of Lycaenidae. Each ROH was classified as either short (< 100 Kb), medium (0.1 - 1 Mb) or long (> 1 Mb) in length.

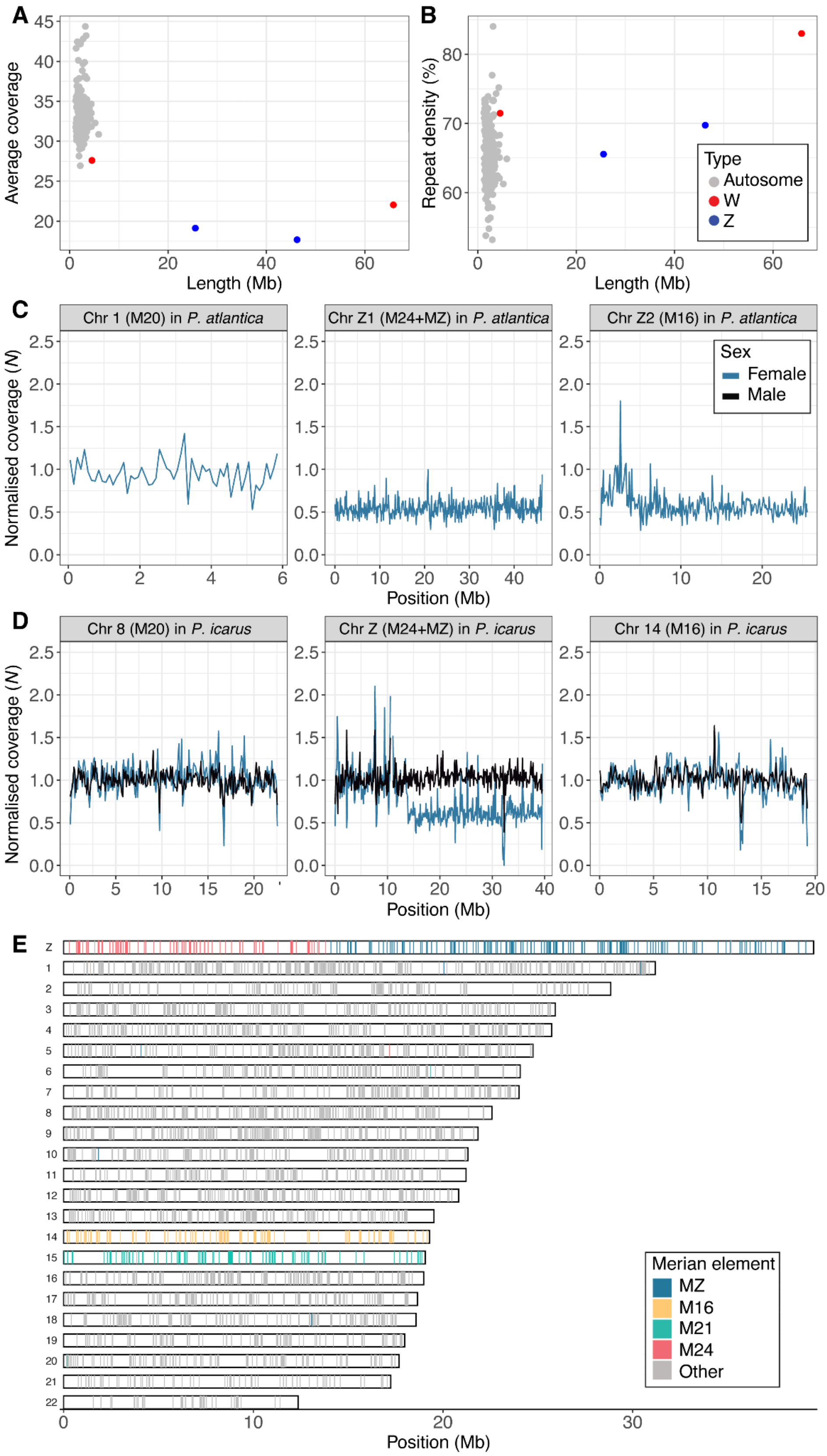

A
Average coverage
Length (Mb)
B
Repeat density (%)
Length (Mb)
Type
Autosome
W
Z
C
Chr 1 (M20) in P. atlantica
Chr Z1 (M24+MZ) in P. atlantica
Chr Z2 (M16) in P. atlantica
Normalised coverage (N)
Sex
Female
Male
Position (Mb)
D
Chr 8 (M20) in P. icarus
Chr Z (M24+MZ) in P. icarus
Chr 14 (M16) in P. icarus
Normalised coverage (N)
Position (Mb)
E
Merian element
MZ
M16
M21
M24
Other
Position (Mb)

**Figure S1. Identification of sex-linked scaffolds in the *P. atlantica* and *P. icarus* reference assemblies reveals *P. atlantica* has two Z chromosomes while *P. icarus* has a single Z chromosome.**

**A.** Average coverage of PacBio data per scaffold where the data was mapped to the final assembly of *P. atlantica.* **B.** Proportion of each scaffold in the *P. atlantica* assembly occupied by repetitive elements. **C.** Normalised female PacBio read coverage in 100 kb windows in the largest autosome (chromosome 1 which corresponds to Merian element 20; M20), Z1 (M24+MZ) and Z2 (M16) of *P. atlantica*. Normalised coverage (*N*) was calculated by dividing the coverage in each 100 kb window by the median autosomal coverage. **D**. Normalised female and male PacBio read coverage in 100 kb windows in chromosome 8 (M20), the Z chromosome (M24+MZ) and chromosome 14 (M16) of *P. icarus.* **E.** Chromosomes of a male *P. icarus* genome, painted with the locations of BUSCO genes. Each gene is coloured by the Merian element that they belong to, if the Merian element is represented on the sex chromosomes of *P. atlantica* (MZ, M16, M21 and M24) or else coloured grey. The Z in *P. icarus* corresponds to the MZ+M24 fusion also found in *P. atlantica*, indicating the fusion was present in the last common ancestor of these two species.

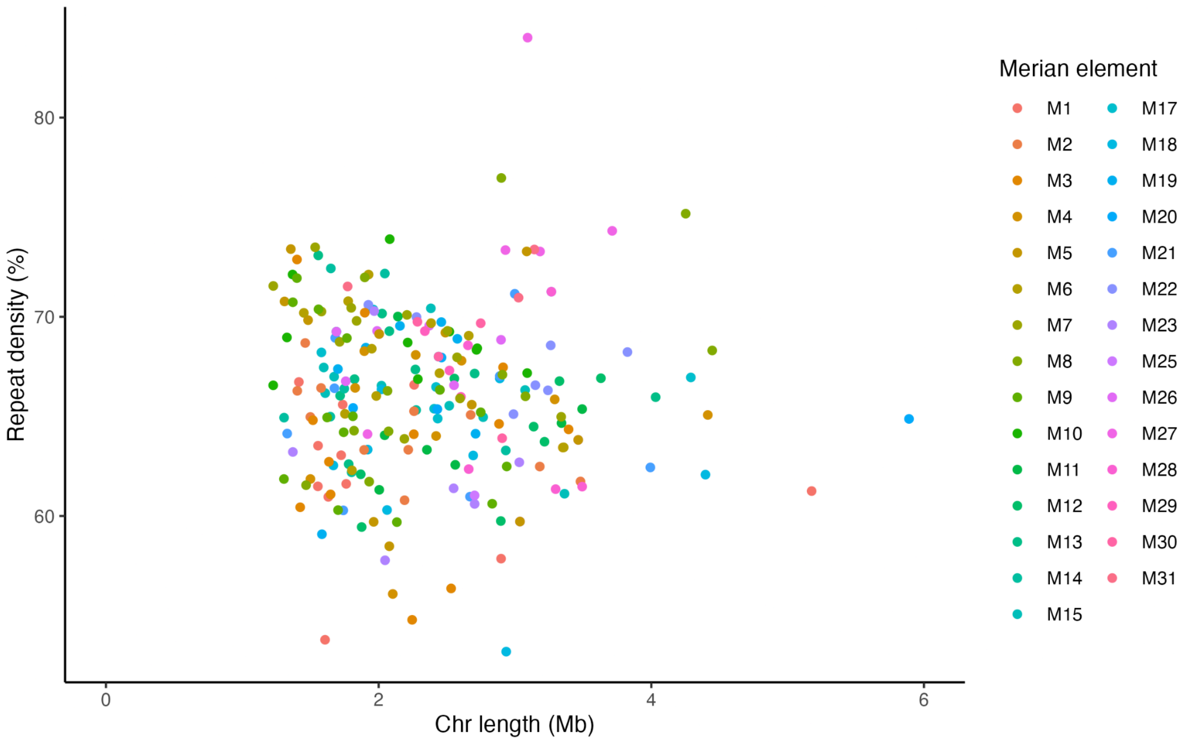


**Figure S2. Autosome length is not correlated with average repeat density in *P. atlantica.***

Chromosome length of each autosome (Mb) against mean repeat density (%) of the autosome. Points are coloured by the Merian element that the chromosome derives from.

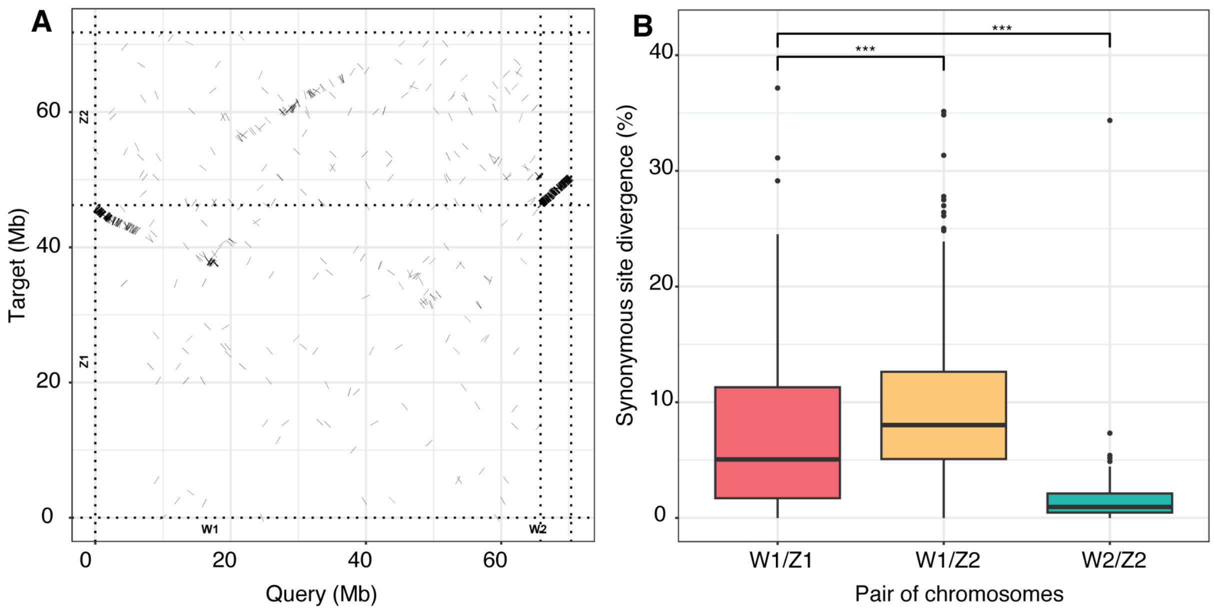


**Figure S3. Nucleotide alignments of W1 and W2 to Z1 and Z2 in *P. atlantica* and distribution of the average synonymous site divergence per pair of sex chromosome**

**A.** Nucleotide alignments between the two Z chromosomes and the two W chromosomes of *P. atlantica.* Alignments were filtered to only retain primary alignments with length >5000 bp and mapping quality >=40. **B.** The average synonymous site divergence between W1 and Z1 is significantly lower than that between W1 and Z2 (Wilcoxon t-test, $p < 0.001$, indicated by ***) and significantly higher than that between W2 and Z2 (Wilcoxon t-test, $p < 0.001$, indicated by ***).

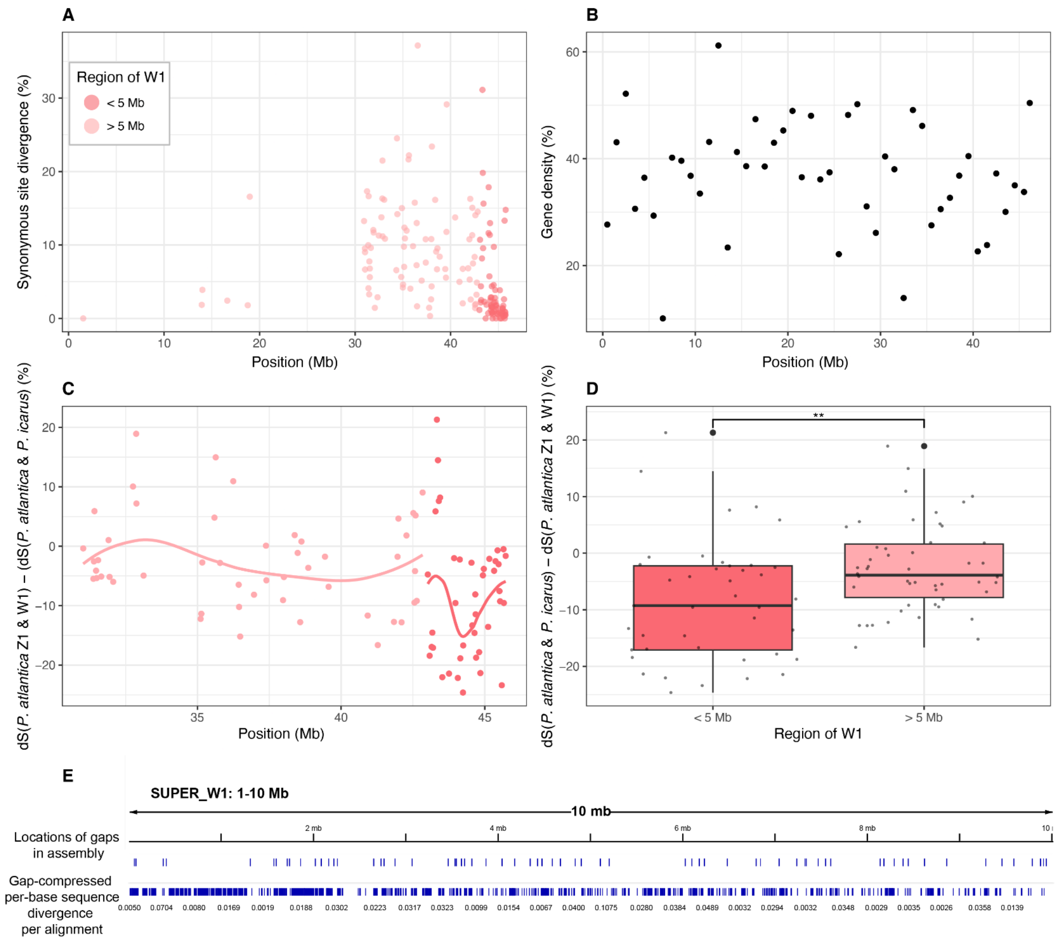


**Figure S4. A region of low synonymous site divergence on Z1 and W1.**

**A.** Synonymous site divergence in 156 orthologues on Z1 and W1 of *P. atlantica* against the position of each orthologue on the Z1 chromosome. Points are coloured by whether the W1 orthologue is located in the first 5 Mb of W1 or not. **B.** Total gene density per 100 kb window along chromosome Z1 of *P. atlantica*. **C.** Distribution of divergence values of Z1 and W1 orthologues in *P. atlantica* normalised by divergence between *P. atlantica* and *P. icarus.* Position of each single-copy protein-coding gene on Z1 that has an orthologue on W1 of *P. atlantica*, against the $d_S$ of the W1 and Z1 orthologues minus the $d_S$ of the Z1 orthologue in *P. atlantica* and *P. icarus*. The position of each orthologue is coloured by whether it is located in the first 5 Mb of W1. The lines represent LOESS smoothing functions fitted to the data. **D.** Boxplot of the values of $d_S$ of the W1 and Z1 orthologues minus the $d_S$ of the Z1 orthologue in *P. atlantica* and *P. icarus*, indicates a significantly lower divergence for orthologue in the first 5 Mb of W1 versus the rest of W1 orthologues with a copy on Z1 (Wilcoxon t-test, $p < 0.01$, indicated by **). Boxplots represent the median and interquartile range, with whiskers set to 1.5 times the interquartile range. **E.** Visualisation of the first 10 kb of the W1 chromosome in *P. atlantica* using IGV annotated with the positions of gaps in the assembly where joins where made by a combination of Hifiasm and during curation, and the gap-compressed per-base sequence divergence of each alignment. This demonstrates that the first 10 kb of W1 includes many gaps, suggesting the low divergence in the 5 Mb is not due to the misassembly of a single region.

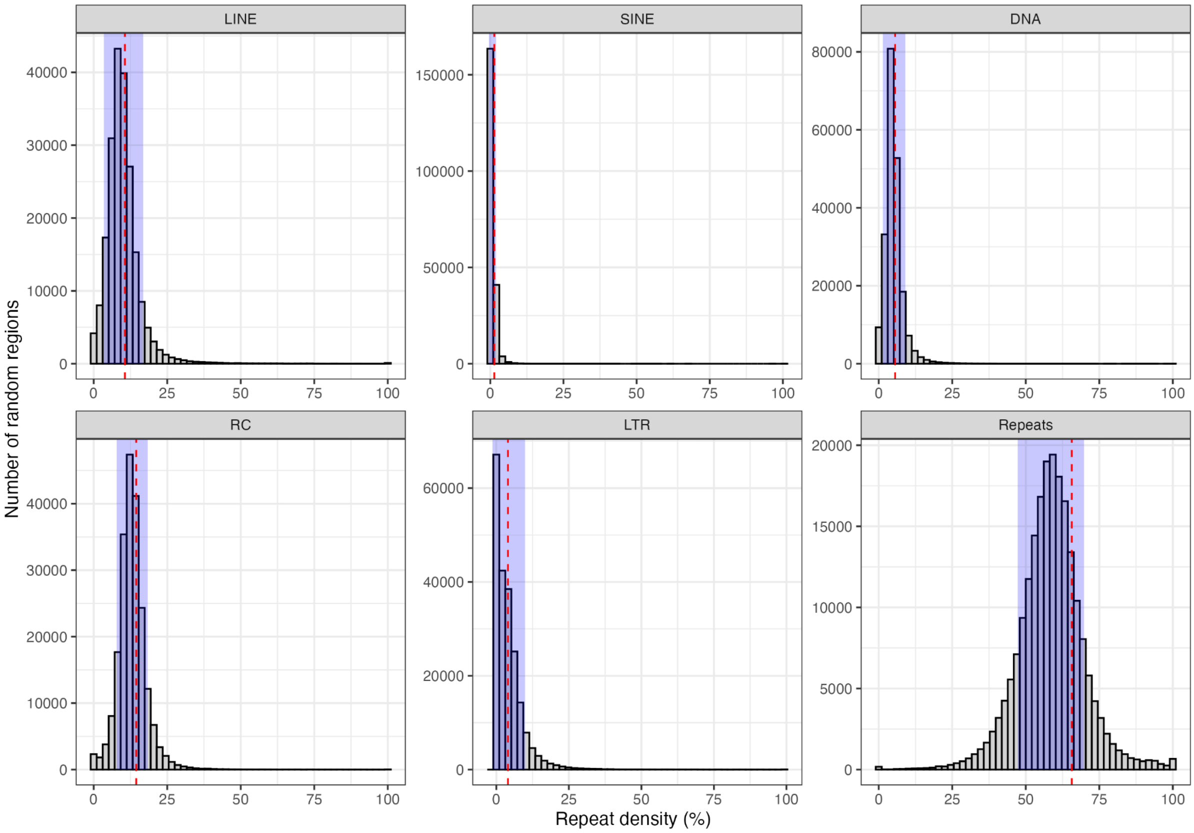


**Figure S5. Fission sites of *P. atlantica* are not associated with highly repetitive regions**

Distribution of density of a given class of transposable elements in 10,000 randomly sampled regions in the *P. icarus* genome (grey bars) compared to the average repeat density observed in regions of the *P. icarus* genome that correspond to regions that underwent fission in *P. atlantica* (red line). The region shaded blue represents the standard deviation in repeat density in the randomly sampled regions. Classes shown are llong interspersed elements (LINEs), short interspersed elements (SINEs), DNA elements (DNA), rolling circle elements (RC), long terminal repeats (LTR), and ‘Other’. The category ‘Other’ includes satellites, simple repeats and low complexity repeats.

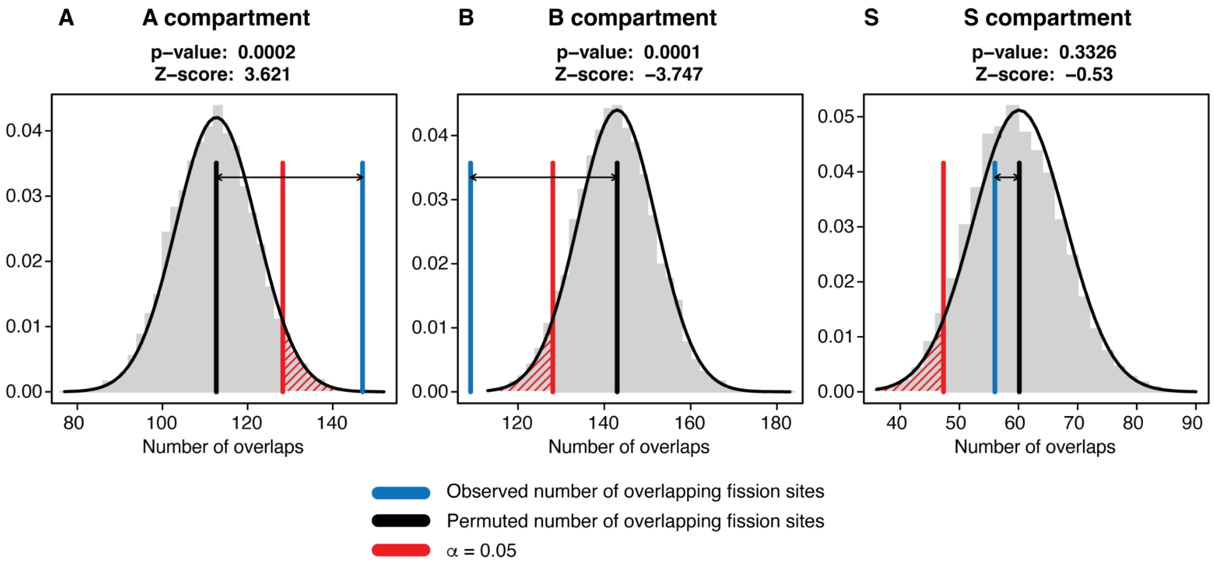


**Figure S6. Correlation between the ancestral chromatin landscape and fission sites in *P. atlantica.***

**A.** Fission sites in *P. atlantica* significantly overlapped with regions of the A compartment of *Cyaniris semiargus* (147/212, (p-value < 0.001, z-score = 3.621). **B.** Fission sites in *P. atlantica* were significantly depleted in the B compartment of *C. semiargus* (p-value < 0.001, z-score = -3.747). **C.** There was no significant overlap between fission sites in *P. atlantica* and the S compartment of *C. semiargus.* The locations of compartments in *C. semiargus* can be used as a proxy for the locations of the compartment in the ancestor of *Polyommatus* before fission occurred. The significance of overlap between fission locations and compartments was assessed with 10,000 resamplings.

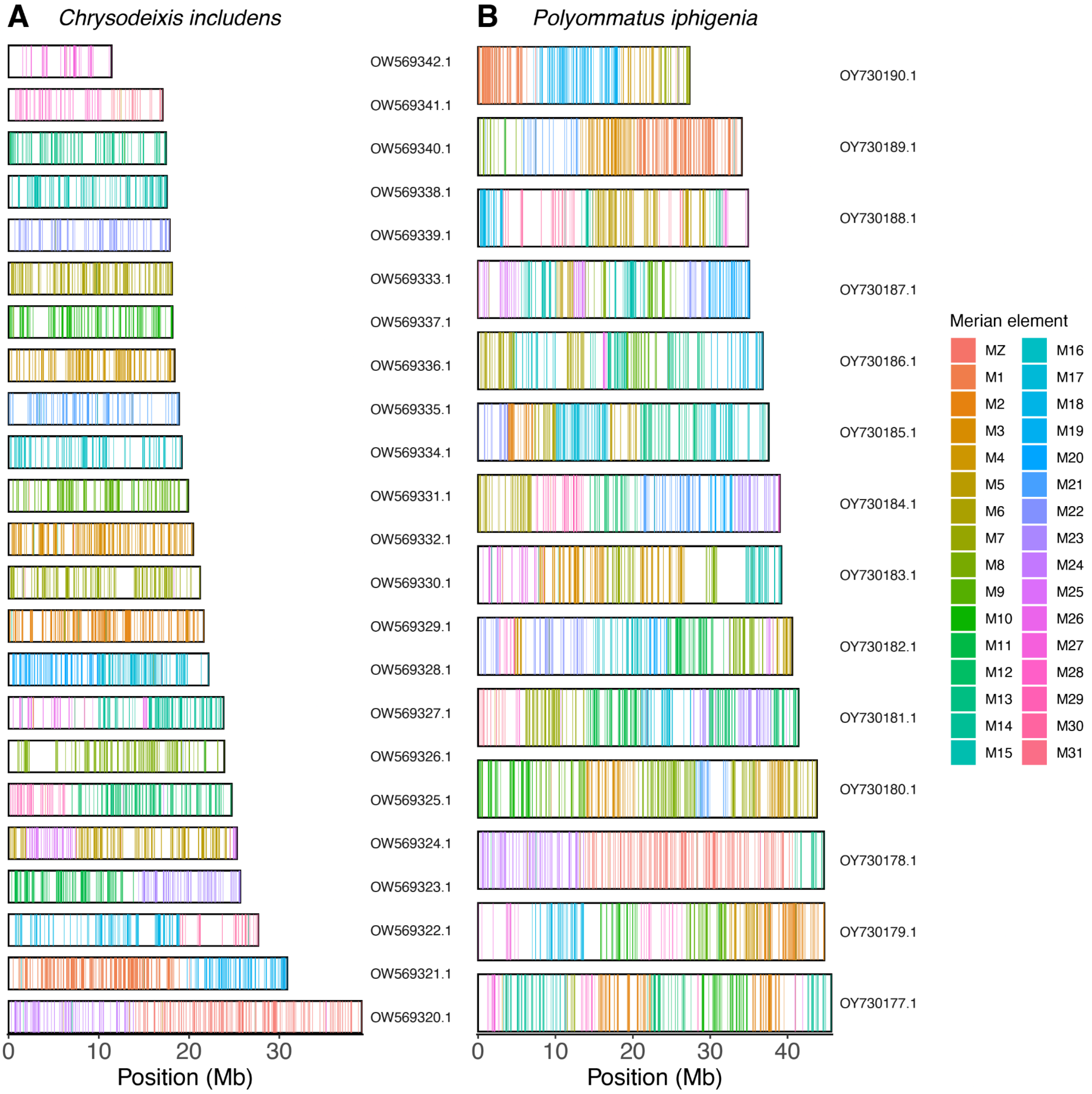


**Figure S7. Merian element paints of *Polyommatus iphigenia* and *Chrysodeixis includens*.**
Chromosomes of **A.** *Chrysodeixis includens* and **B.** *Polyommatus iphigenia.* Each chromosome is represented by a rectangle and the positions of genes are indicated by lines. Lines are coloured by the Merian element that the gene belongs to.

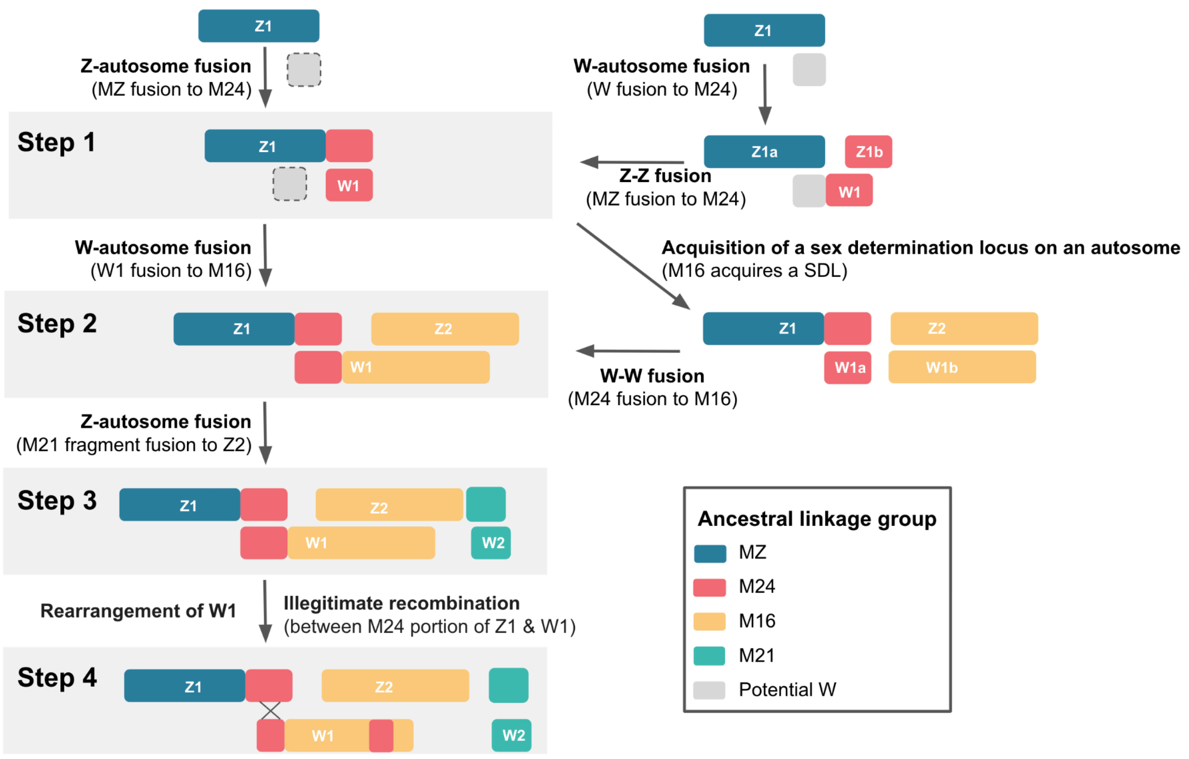


**Figure S8. Models of potential routes by which the Z1Z2/W1W2 chromosome system in *P. atlantica* evolved.**

We propose that the sex chromosomes of *P. atlantica* arose through four key steps. Each step corresponds to an intermediate state supported by the data. However, there are multiple routes by which each intermediate state may have been reached.

Chromosomes are coloured by the ancestral linkage group that the chromosome evolved from. The sex chromosomes in the ancestor of *P. atlantica* was either Z0 or ZW before additional sex chromosomes evolved.

**Step 1:** The autosomal Merian element (M24) fused to either the ancestral heterogametic Z chromosome (M24+MZ) or to a W chromosome (M24+W). If a W-autosome fusion occurred, the homolog of the autosome that fused to the W chromosome was initially a separate chromosome but then underwent fusion with the ancestral Z chromosome resulting in the observed single large Z chromosome (MZ+M24).

**Step 2:** An autosomal Merian element (M16) became a neo-Z chromosome, causing the autosome to become sex linked. This occurred in one of two ways: either M16 fused to W1 or M16 first acquired a locus associated with sex determination. Under the second scenario, one M16 subsequently fused to W1 (W1a+W1b) to form a single larger W chromosome.

**Step 3:** An autosomal fragment corresponding to a portion of Merian element 21 (M21) fused to Z2. The homolog of this chromosome would therefore have become a second W chromosome (W2). A Z-autosome fusion is more likely than a W-autosome fusion in this step as M21-derived genes are not located on W2.

**Step 4:** The W chromosome underwent intrachromosomal rearrangement. A rare recombination event may have occurred between the M24-derived sequence of Z1 and W1.

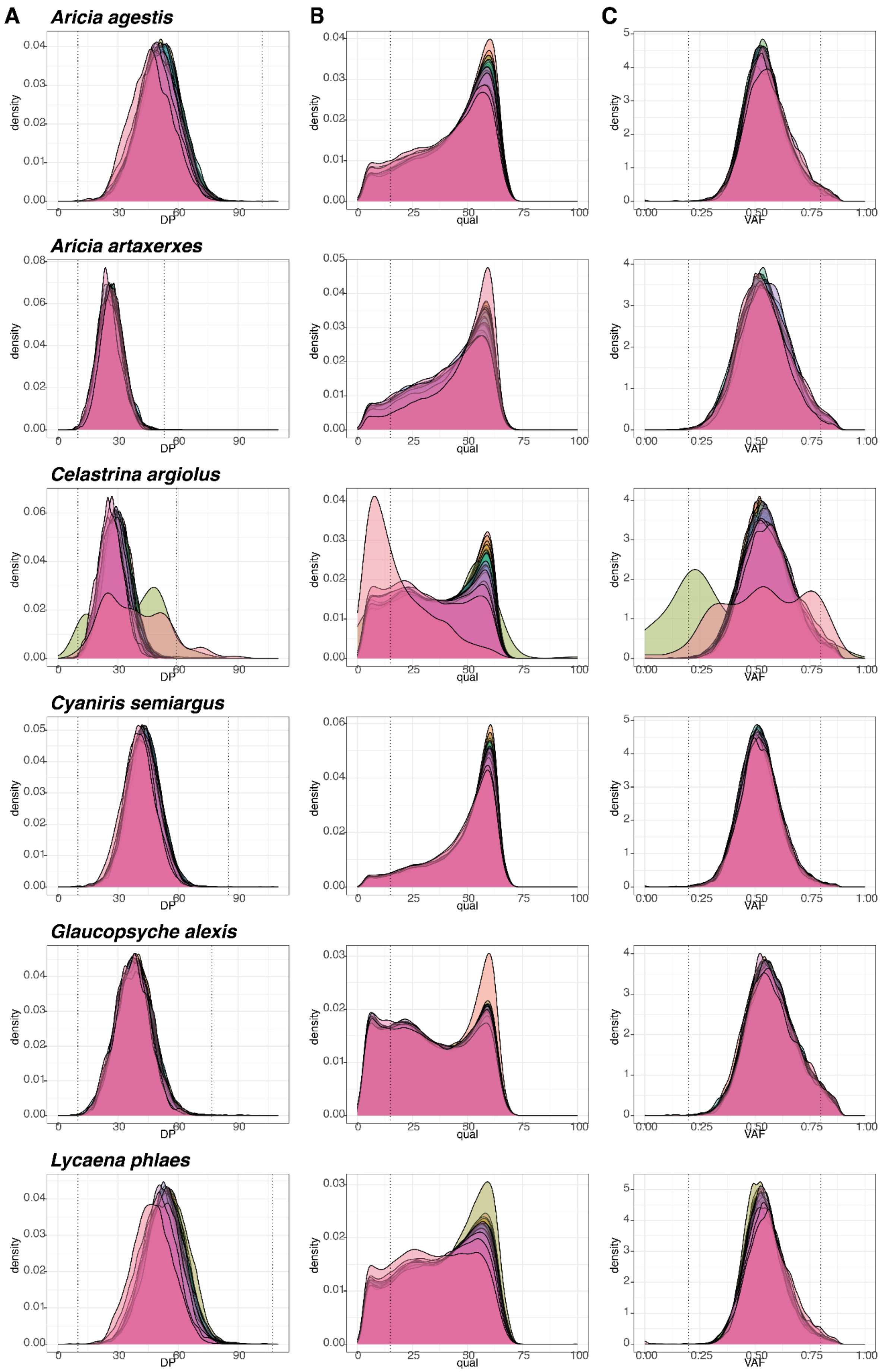
A
B
C
Aricia agestis
Aricia artaxerxes
Celastrina argiolus
Cyaniris semiargus
Glaucopsyche alexis
Lycaena phlaes
density
DP
qual
VAF

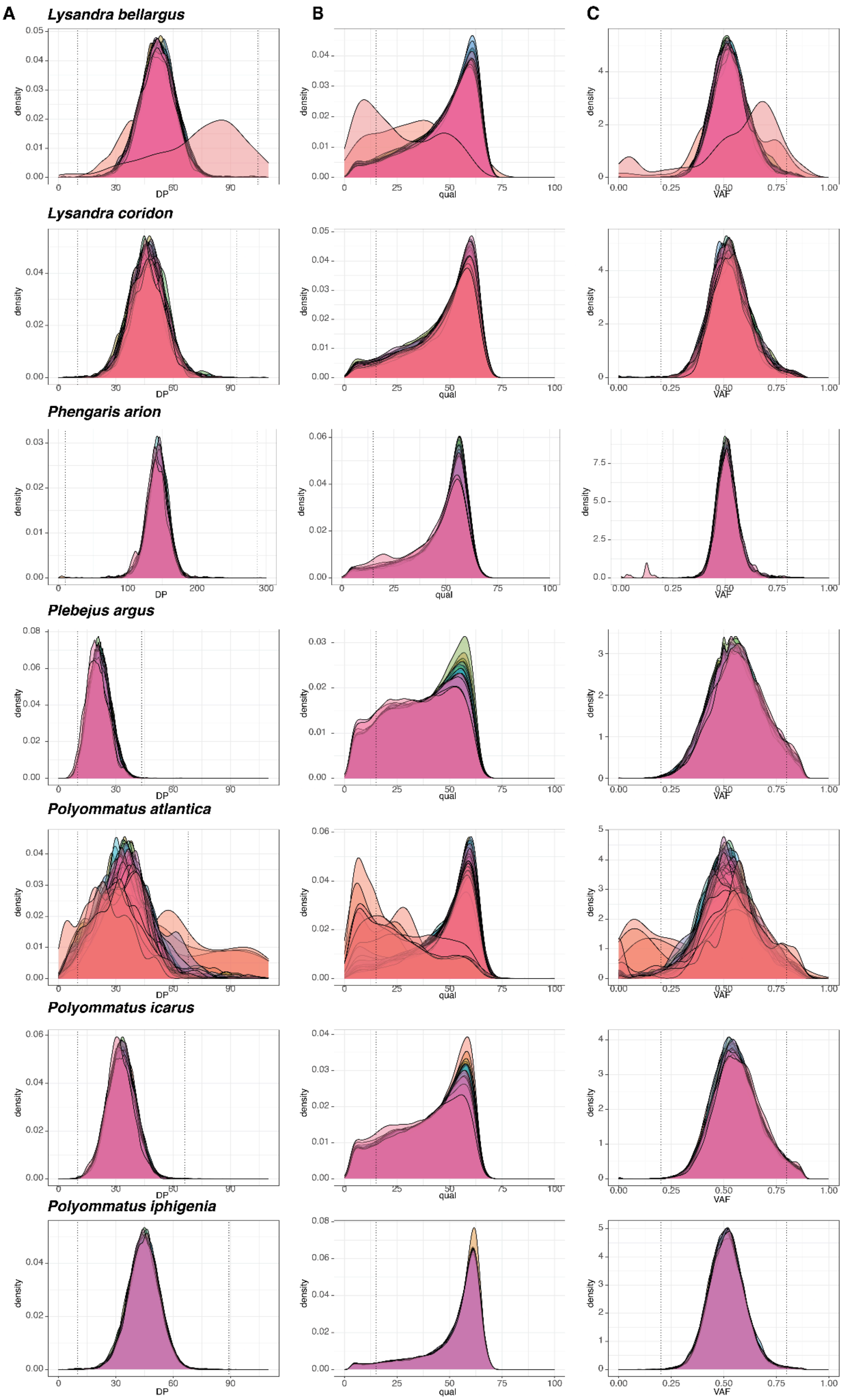
A
B
C
Lysandra bellargus
Lysandra coridon
Phengaris arion
Plebejus argus
Polyommatus atlantica
Polyommatus icarus
Polyommatus iphigenia
density
DP
qual
VAF

**Figure S9. Quality control and filtering of variants.**

Quality control of variant calling and selection of filtering criteria. Distribution of **A.** depth, **B.** PHRED_quality scores (QUAL) and **C.** variant allele frequency (VAF) for each SNP. Distributions are plotted per species. Grey lines indicate (A) a depth of 10 and twice the mean depth per genome, (B) a minimum QUAL of 15, (C) VAF of 0.2 and 0.8.

1. Hayward, A., Lohse, K., Vila, R., Laetsch, D.R., Hedlund, J.S.U., Wellcome Sanger Institute Tree of Life programme, Wellcome Sanger Institute Scientific Operations: DNA Pipelines collective, Tree of Life Core Informatics collective, and Darwin Tree of Life Consortium (2023). The genome sequence of the Brown Argus, *Aricia agestis* (Denis & Schiffermüller, 1775). Wellcome Open Res. *8*, 336.

2. Ebdon, S., Lohse, K., Jansen Van Rensburg, A., Wellcome Sanger Institute Tree of Life programme, Wellcome Sanger Institute Scientific Operations: DNA Pipelines collective, Tree of Life Core Informatics collective, and Darwin Tree of Life Consortium (2022). The genome sequence of the northern brown argus, Aricia artaxerxes (Fabricius, 1793). Wellcome Open Res. *7*, 314.

3. Hayward, A., Wright, C., Darwin Tree of Life Barcoding collective, Wellcome Sanger Institute Tree of Life programme, Wellcome Sanger Institute Scientific Operations: DNA Pipelines collective, Tree of Life Core Informatics collective, and Darwin Tree of Life Consortium (2021). The genome sequence of the holly blue, Celastrina argiolus (Linnaeus, 1758). Wellcome Open Res *6*, 340.

4. Lohse, K., Hayward, A., Laetsch, D.R., Marques, V., Vila, R., Tyler-Smith, C., Wellcome Sanger Institute Tree of Life programme, Wellcome Sanger Institute Scientific Operations: DNA Pipelines collective, Tree of Life Core Informatics collective, and Darwin Tree of Life Consortium (2023). The genome sequence of the Mazarine Blue, Cyaniris semiargus (Rottemburg, 1775). Wellcome Open Res. *8*, 181.

5. Hinojosa Galisteo, J.C., Vila, R., Darwin Tree of Life Barcoding collective, Wellcome Sanger Institute Tree of Life programme, Wellcome Sanger Institute Scientific Operations: DNA Pipelines collective, Tree of Life Core Informatics collective, and Darwin Tree of Life Consortium (2021). The genome sequence of the green-underside blue, Glaucopsyche alexis (Poda, 1761). Wellcome Open Res. *6*, 274.

6. Lohse, K., Laetsch, D.R., Vila, R., Darwin Tree of Life Barcoding collective, Wellcome Sanger Institute Tree of Life programme, Wellcome Sanger Institute Scientific Operations: DNA Pipelines collective, Tree of Life Core Informatics collective, and Darwin Tree of Life Consortium (2021). The genome sequence of the small copper, Lycaena phlaeas (Linnaeus, 1760). Wellcome Open Res. *6*, 294.

7. Lohse, K., Hayward, A., Vila, R., Howe, C., Wellcome Sanger Institute Tree of Life programme, Wellcome Sanger Institute Scientific Operations: DNA Pipelines collective, Tree of Life Core Informatics collective, and Darwin Tree of Life Consortium (2022). The genome sequence of the Adonis blue, Lysandra bellargus (Rottemburg, 1775). Wellcome Open Res. *7*, 255.

8. Vila, R., Lohse, K., Hayward, A., Laetsch, D.R., Wright, C., Wellcome Sanger Institute Tree of Life programme, Wellcome Sanger Institute Scientific Operations: DNA Pipelines collective, Tree of Life Core Informatics collective, and Darwin Tree of Life Consortium (2023). The genome sequence of the Chalkhill Blue, Lysandra coridon (Poda, 1761). Wellcome Open Res. *8*, 162.

9. Meredith, S.A., Simcox, D.J., Thomas, J.A., Sumnall, A., Holland, P.W.H., Crowley, L.M., University of Oxford and Wytham Woods Genome Acquisition Lab, Darwin Tree of Life Barcoding collective, Wellcome Sanger Institute Tree of Life Management, Samples and Laboratory team, Wellcome Sanger Institute Scientific Operations: Sequencing Operations, et al. (2024). The genome sequence of the Large Blue butterfly, Phengaris (Maculinea) arion (Linnaeus, 1758). Wellcome Open Res. *9*, 506.

10. Hayward, A., Lohse, K., Laetsch, D.R., Vila, R., Wellcome Sanger Institute Tree of Life programme, Wellcome Sanger Institute Scientific Operations: DNA Pipelines collective, Tree of Life Core Informatics collective, Taluy, E., and Darwin Tree of Life Consortium (2022). The genome sequence of the silver-studded blue, Plebejus argus (Linnaeus, 1758). Wellcome Open Res *7*, 315.

11. Lohse, K., Wellcome Sanger Institute Tree of Life programme, Wellcome Sanger Institute Scientific Operations: DNA Pipelines collective, Tree of Life Core Informatics collective, and

Darwin Tree of Life Consortium (2023). The genome sequence of the Common Blue, Polyommatus icarus (Rottemburg, 1775) [version 1; peer review: awaiting peer review]. Wellcome Open Res. *8*, 72.
